# Combinatorial mechanisms specify cellular location and neurotransmitter identity during regeneration of planarian neurons

**DOI:** 10.1101/2025.05.23.655781

**Authors:** Kendall B. Clay, Taylor Medlock-Lanier, Rachel N. Grimes, Olabamibo O. Oke, Nikolay M. Filipov, Rachel H. Roberts-Galbraith

**Affiliations:** Neuroscience Program, University of Georgia, Athens, GA 30602, USA; Department of Cellular Biology, University of Georgia, Athens, GA 30602, USA; Department of Biochemistry and Molecular Biology, University of Georgia, Athens, GA 30602, USA; Department of Physiology and Pharmacology, University of Georgia, Athens, GA 30602, USA; Interdisciplinary Toxicology Program, University of Georgia, Athens, GA, 30602, USA

**Keywords:** regeneration, dopamine, nervous system, planarian, regenerative neurogenesis, neuronal cell fate, specification, transcriptional regulation, differentiation

## Abstract

During regenerative neurogenesis, neurons must be created in the right types and locations. Though regenerative neurogenesis is limited in humans, other animals use regenerative neurogenesis to faithfully restore form and function after brain injury. Planarians are flatworms with extraordinary capacity for brain regeneration. Planarians use pluripotent stem cells to create neurons after injury, rather than resident progenitors. In this context, genetic mechanisms that produce diverse neurons with correct local identity remain unknown. Here, we report the discovery of factors important for regenerative neurogenesis of dopaminergic neurons in the planarian central, peripheral, and pharyngeal nervous systems. Distinct genes promote dopaminergic neuronal identity and instruct neurons to inhabit regions of the nervous system. Our results demonstrate that planarian neuronal fate requires factors that simultaneously direct neurotransmitter choice and regional location. Our work suggests that combinatorial direction of cell type could inform and improve exogenous stem cell therapies aimed at precisely replacing neurons.

## Introduction

Developmental neurogenesis involves neuronal birth from progenitors with predictable timing, relatively fixed birth order, and within specific locations within a growing animal. In contrast, regenerative neurogenesis occurs outside normal developmental timing, with nascent cells and surviving cells intermingling unpredictably at the injury site. Repairing injury also requires an unpredictable number and type of neurons, with the ideal outcome being faithful reproduction of correct cell types and ratios for a given location. In some organisms, regenerative neurogenesis occurs through a pool of localized progenitors that give rise to mature cell types in distinct regions^1^. However, in many animals including humans, regenerative neurogenesis in the central nervous system (CNS) occurs infrequently in only a few areas^2,3^. Therefore, understanding how to induce regenerative neurogenesis and how to introduce exogenous stem cells directed toward injury-specific regenerative outcomes could revolutionize therapies for neurodegenerative diseases and other CNS injuries.

We are interested in developing a model system with which to understand successful regenerative neurogenesis. Planarian flatworms are known for their regenerative abilities, including *de novo* brain regrowth after nearly any injury^4–6^. In planarians, pluripotent stem cells respond to injury by dividing and generating progeny^7–10^. Critically, neurons are produced in the correct numbers, diversity, pattern, and connectivity to restore function^4,11^. The ability of planarians to regenerate neurons precisely without resident progenitors raises many interesting questions: How do pluripotent stem cells establish neurons in the right numbers, types, and ratios? How are neuronal cell identity and localization specified during regenerative neurogenesis? These questions merit answers because human stem cell-based therapies will likely rely on exogenous stem cells, with the requirement that these cells accurately differentiate, mature, localize, and connect to restore function to damaged neural tissue.

In addition to regenerative capacity, the planarian nervous system has spatial and cell type complexity, making it suitable for study. The planarian nervous system is composed of three parts: the CNS that includes brain and ventral nerve cords; the peripheral nervous system (PNS); and a pharyngeal nervous system (PhNS)^5,12,13^. The planarian nervous system includes at least 70 neuronal cell types^5^, with each cell type present in predictable numbers and ratios^11,14–16^, and glial cells^17,18^. All cell types are replaced after injury *and* are replenished during animal growth and homeostasis^19^. Some regulators of neuronal and glial identities have been identified^5,15,26–35,17,36,37,18,20–25^. For example, *pitx* and *lhx1/5-1* direct terminal differentiation of serotoninergic neurons^25^. And *soxB1-2*, *coe,* and *sim* play broad roles in neuronal specification^5,23,24^. However, the full pathway from pluripotent stem cell to mature neuron has not been determined for any cell type in the planarian nervous system.

To better understand how regenerative neurogenesis proceeds in planarians, we focused on a single, conserved cell type: dopaminergic neurons. Across metazoans, dopamine is synthesized through conserved enzymes tyrosine hydroxylase (TH) and amino acid decarboxylase (AADC)^38^. In planarians, *TH* and *AADC*^39^ mark a subset of neurons that are easily quantifiable *and* present in all areas of the nervous system. Additionally, *TH^+^* neurons have important functions in locomotion in planarians and other animals. Similar phenotypes arise through pharmacological modulation of dopamine receptors, tracing these functions to dopamine specifically^39,40,49,50,41–48^. Translationally, loss of dopaminergic neurons is central to Parkinson’s Disease^51–53^ and implicated in other neurodegenerative diseases^54–57^. Both experimental tractability and deep conservation made dopaminergic neurons a well-suited starting point for understanding mechanisms directing neurogenesis in the context of planarian regeneration.

Here, we sought to understand the molecular basis of dopaminergic regeneration in planarians. We completed a reverse genetic screen targeting 74 candidate genes enriched in planarian *TH^+^* neurons. We identified 11 genes important for dopaminergic neuron regeneration and maintenance. Interestingly, we found that *friend leukemia integration 2* (*fli1-2*) and *iroquois 4/6* (*irx-4/6*) regulate dopaminergic neurons body-wide, while 9 additional genes promote dopaminergic neurons in a region-specific manner. For example, *lim only domain 1/3-1* (*lmo1/3-1*) impacts regeneration of *TH^+^* cells in the brain alone. We also found that one gene, *amyloid precursor protein-like 1 (app-L1)*, is critical for *TH*^+^ neurons of the brain in a regeneration-specific context. Altogether, our data support a model in which a core group of transcription factors works cell-autonomously to drive dopaminergic neuron fate and maturation, while distinct factors drive tissue- and region-specificity. We conclude that combinatorial codes work together to guide dopaminergic neuronal production for each region of the nervous system.

## Results

### A subset of genes co-expressed with TH is required for regeneration of TH^+^ cells

Planarians have a complex nervous system, with central, peripheral, and pharyngeal networks of cells^12,13^ (Fig. 1A). Markers of dopaminergic neurons have been reported, including genes encoding two enzymes involved in the biosynthesis pathway, *TH* and *AADCa*^39^. We validated these markers in *Schmidtea mediterranea*, our model planarian (Fig. 1B). We wanted an additional marker independent of the dopamine biosynthesis pathway. *slc6a3*^58^, a homolog of the dopamine transporter *dat*, is also nearly always co-expressed with *TH* (Fig. 1B-C). In the literature, *TH*^+^ cells in planarians have been referred to as dopaminergic neurons. However, in other organisms, norepinephrine is synthesized from dopamine by dopamine beta hydroxylase (DBH) or alternatively by tyramine beta hydroxylase (TBH)^59–63^. In planarians, only one *TBH/DBH* homolog has been identified^36,64,65^. Though *TH^+^*and *TBH*^+^ staining was previously determined to be nonoverlapping in *Dugesia japonica*^65^, we find a rare population of *TH^+^/TBH*^+^ neurons that are potentially norepinephrinergic (Fig. 1D). Supporting this finding, we showed that *TH*(RNAi) resulted in a dramatic decrease of both dopamine and norepinephrine, measured by high performance liquid chromatography (HPLC) (Fig. 1E). We do note that the vast majority (95%) of *TH^+^* neurons in the brain are *TPH^-^* and *<u>all</u> TH^+^* neurons in the PNS and PhNS are *TH^+^*/*TPH^-^.* These neurons are likely all dopaminergic. Importantly, we noted no co-expression of *slc6a3* and *TBH* (Fig. 1D), so norepinephrine is likely packaged into vesicles by another transporter and *slc6a3* can be used as a marker of dopaminergic neurons that do not produce norepinephrine.

**Figure 1:**
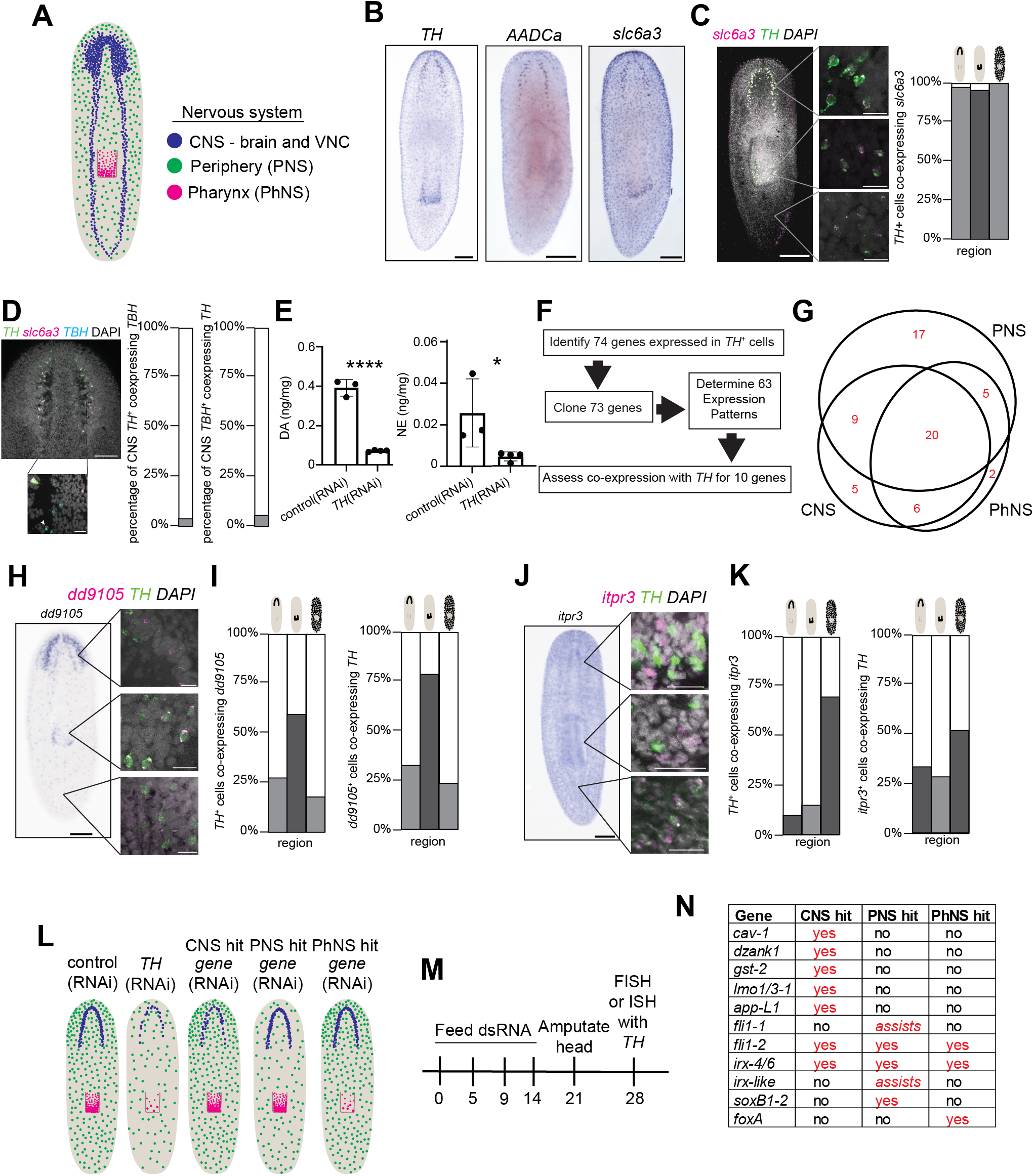
A transcriptomic approach to identifying genes responsible for the regeneration of dopaminergic neurons in planarians. A) Schematic of the planarian nervous system showing the central (CNS) in blue, periphery (PNS) in green, and pharyngeal (PhNS) in magenta. B) Whole-body ISH showing markers for dopaminergic neurons: *TH*, *AADCa*, and *slc6a3*. C) Co-expression of *TH* and *slc6a3* with quantification. D) Co-expression of *TH* and *TBH* with quantification. E) Amount of dopamine and norepinephrine detected in control and *TH* knockdown samples. F) Screen workflow for determining expression patterns and assessing co-expression with *TH.* G) Euler diagram showing where the 64 genes are expressed – areas are proportional to the number of candidates in that category. Overlap indicates a gene is expressed in both regions. H-K) Representative images and quantification of co-expression with *TH* for two genes (*dd9105* and *itpr3*) found in all three regions of the nervous system. Schematics above the bars indicate the region of the nervous system for quantitation. For each gene, a colorimetric ISH is shown on the left, with insets showing co-expression of the gene of interest (magenta), *TH* (green), and DAPI (nuclei – gray) visualized after FISH. The percentage of *TH*^+^ co-expression is shown in the tiara, pharynx, and PNS in both directions. L) Schematic showing expected results from control(RNAi), *TH*(RNAi), and potential hits in the screen that impact distinct regions of the nervous system. M) RNAi paradigm used in our RNAi screen. N) Table of genes for which RNAi caused a reduction of dopaminergic neurons in one or more regions of the nervous system. (* P < 0.05, **** P < 0.0001) Scale bars = 200 µm (B, C, I, J except insets), 50 µm (D except inset), 20 µm (C, D, I, J insets).

After better characterizing dopaminergic neurons in planarians, we next sought to identify genes important for their regeneration and maintenance. We searched single-cell transcriptomic data, identifying 74 genes with high fold enrichment in cell clusters enriched in transcripts for *TH*, *AADC*, and *slc6a3*^14^ (Fig. 1F). To confirm neuronal expression of these genes, we cloned 73/74 transcripts and examined expression patterns for 64 of these genes. We determined that all transcripts examined (64/64) were expressed in the CNS, PNS, and/or PhNS (Fig. 1G, Extended Fig. 1A-H). A minority of genes (7/64) were also expressed in cells outside of the nervous system (Extended Fig. 1H). To confirm relevance to dopaminergic neurons, we quantified co-expression of *TH* and 10 representative transcripts. We saw significant heterogeneity in the amount and spatial specificity of co-expression. However, all genes studied were co-expressed with *TH*, validating our discovery strategy (Fig 1H-K, Extended Fig. 2A-P).

Finally, we tested whether each candidate gene plays a role in regeneration of dopaminergic neurons. We knocked down each gene through RNA interference (RNAi) by feeding double stranded RNA (dsRNA) and quantified *TH*^+^ cells after regeneration using colorimetric *in situ* hybridization (ISH) and/or fluorescent *in situ* hybridization (FISH) (Fig. 1L-M). We identified 11 genes required for regeneration and/or maintenance of dopaminergic neurons in one or more parts of the nervous system through this strategy (Fig. 1N and subsequent figures). Separately, we validated knockdown efficiency and specificity of our RNAi experiments using RT-qPCR (Extended Fig. 3).

### lmo1/3-1 and app-L1 play specific roles in the regeneration of CNS dopaminergic neurons

To confirm the results of our primary screen, we repeated RNAi of 15 genes and performed FISH to quantify *TH*^+^ cells in the brain. Because planarian cell number scales readily with body area^66,67^, we normalized brain *TH*^+^ cell number to body size. We determined that RNAi targeting five genes caused fewer *TH*^+^ cells to form in the regenerating brain: *caveolin-1 (cav-1), double zinc ribbon and ankyrin repeat domains 1 (dzank1), glutathione S-transferase-2 (gst-2), lmo1/3-1*, and *app-L1* (Fig. 2A-B). We validated the decrease in dopaminergic neuron numbers by quantifying *slc6a3^+^*cells (Extended Fig. 4A-B).

**Figure 2:**
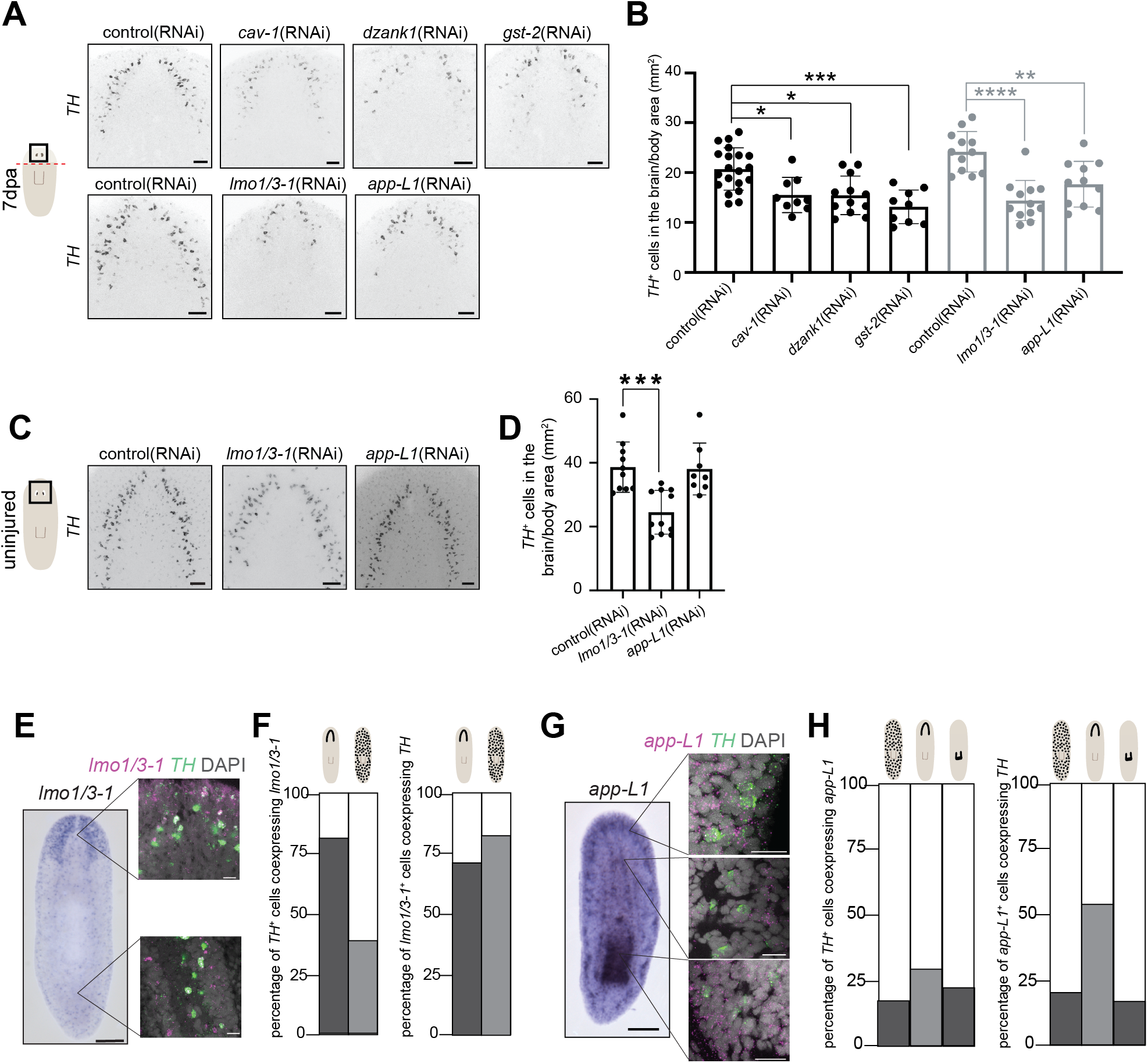
*lmo1/3-1* and *app-L1* are required for the proper regeneration of dopaminergic neurons in the CNS. A) Inverted grayscale FISH images showing *TH^+^* cells in the regenerated tiara 7 days post amputation (dpa) after knockdown of either *cav-1, dzank1, gst-2, lmo1/3-1* or *app-L1* compared to control. B) Quantification of A. One-way ANOVA with Dunnett’s multiple comparisons test. C) Inverted grayscale FISH images showing *TH*^+^ cells in the tiara in uninjured animals after knockdown of *lmo1/3-1* or *app-L1* compared to control. D) Quantification of C. One-way ANOVA with Dunnett’s multiple comparisons test. E-H) Representative images and quantification of co-expression of *lmo1/3-1* and *app-L1* with *TH*. Schematics above the bars indicate the region of the nervous system. For each gene, a colorimetric ISH is shown on the left, with insets showing co-expression of the gene of interest (magenta), *TH* (green), and DAPI (nuclei – gray) visualized after FISH. The percentage of *TH*^+^ cells co-expressing each gene is shown in the CNS and PNS for both and the PhNS for *app-L1*. (* P < 0.05, ** P < 0.01, *** P < 0.001, **** P < 0.0001). Scale bars = 200 µm (E and G except insets), 50 µm (A, C), 20 µm (E and G insets).

We chose to follow up on two of these genes. We prioritized study of *lmo1/3-1* because *lmo3* has been implicated in dopaminergic neuron specification in other models^68–70^ and because LMO proteins often function as part of LIM-domain binding protein transcriptional complexes, which our lab previously investigated in planarians^71^. Additionally, we chose *app-L1* (for phylogenetic explanation of nomenclature, see Supp. File 1) due to physiological roles for APP products in neural development^72^ and due to the connection between APP-derived peptide aggregation and neurodegeneration^73–75^ (for review, see^76–78^).

Because we saw roles for *lmo1/3-1* and *app-L1* in regeneration of *TH*^+^ cells of the brain, we next investigated whether they promoted maintenance of dopaminergic neurons. In this context, we saw a significant reduction of *TH*^+^ cells after *lmo1/3-1*(RNAi) but no change after *app-L1*(RNAi) in animals subjected to RNAi but not amputation (Fig. 2C-D). To rule out a role for *app-L1* in maintenance of dopaminergic neurons, we repeated the experiment with an even longer paradigm (Extended Fig. 4C). The resulting worms still had typical *TH*^+^/*slc6a3*^+^ cell number, confirming *app-L1*’s regeneration-specific role on dopaminergic neurons (Extended Fig. 4D-F).

To confirm that *lmo1/3-1* and *app-L1* could work cell-autonomously, we completed co-expression analysis with *TH* and either *lmo1/3-1* or *app-L1*. Consistent with prior findings, *lmo1/3-1* is expressed in the brain and PNS, frequently in *TH^+^* cells (Fig. 2E-F)^71^. *app-L1* is expressed broadly in the head, pharynx, epidermis, and protonephridia through our ISH (Fig. 2G) and in single-cell sequencing data^14,16^, and is co-expressed with *TH*^14^ (Fig. 2G-H). Based on these analyses, we conclude that *lmo1/3-1* and *app-L1* could impact *TH^+^* neurons cell-autonomously.

Because *lmo1/3-1* and *app-L1* are co-expressed with *TH* in the PNS, we next asked whether *lmo1/3-1* and *app-L1* regulate peripheral dopaminergic neurons. After assessing *TH*^+^ cells in regenerated and maintained tails, we confirmed there was no change in peripheral *TH*^+^ cell number after *lmo1/3-1*(RNAi) or *app-L1*(RNAi) (Extended Fig. 5A-D). *TH^+^* cells in the pharynx express *app-L1* but not *lmo1/3-1* (Fig. 2E-H), so we tested whether *app-L1* is required for pharyngeal dopaminergic neurons. We observed no change in pharyngeal *TH^+^*cells following *app-L1*(RNAi) in regeneration or maintenance (Extended Fig. 5E-H).

We then investigated the specificity of *lmo1/3-1* or *app-L1* phenotypes. We first looked at a broad marker of the CNS, *choline acetyl transferase* (*ChAT*)^79^ after RNAi. Regenerated brain size, normalized to body size, was unchanged after *lmo1/3-1*(RNAi), *app-L1*(RNAi), or *TH*(RNAi) (Extended Fig. 6A-B). Because *lmo1/3-1* is expressed in some *TH^-^* neurons in transcriptional atlases (e.g. *cpp-1^+^* peptidergic neurons), we examined the impact of *lmo1/3-1*(RNAi) on *cpp-1^+^* cells^14,16,22^. *lmo1/3-1*(RNAi) caused a significant reduction in *cpp-1^+^* neuron*s* after head regeneration, indicating that *lmo1/3-1* directs neuronal regeneration for a limited number of additional neuron types (Extended Fig. 6C-D).

Additionally, due to our discovery that some *TH*^+^ cells express *TBH*, we wanted to assess octopaminergic (*TBH*^+^/*TH*^-^) and norepinephrinergic neurons (*TBH*^+^/*TH*^+^) after loss of *lmo1/3-1* or *app-L1*. We found that *lmo1/3-1*(RNAi) led to a significant reduction in dopaminergic, octopaminergic, and norepinephrinergic neurons, but *app-L1*(RNAi) reduced dopaminergic neurons specifically (Extended Fig. 6E-H). Due to the high level of co-expression between *lmo1/3-1* and *TH*, we posit that *lmo1/3-1* plays a significant, but not specific, role in dopaminergic neurons in the brain. Despite *app-L1*’s broad expression pattern, its role in the nervous system appears to be narrow, impacting dopaminergic neurons but not neurons in general.

### FLI and IRX genes are required for TH^+^ cells throughout the nervous system

In the original screen, we performed head amputations to assess brain regeneration. Surprisingly, after knockdown of *fli1-2* or *irx-4/6*, we observed a striking reduction of dopaminergic neurons in the PNS (Fig. 3A-B). *fli1-2* is synonymous with previously described *fli1*^36^ and *irx-4/6* corresponds to previously described *irx-2*^15^. Notably, planarian *irx-4/6* had previously been implicated as a fate-specific transcription factor (FSTF) for *TH^+^* neurons^15^. We decided to follow up on *fli1-2* and *irx-4/6,* and their related family members — *fli1-1* (previously *fli-1*^18^ and *elf-1*^36^) and *irx-like —* which were also enriched in *TH^+^* cells. We note that each nomenclature adjustment was based on phylogenetic analyses (Supp. File 2-3).

**Figure 3:**
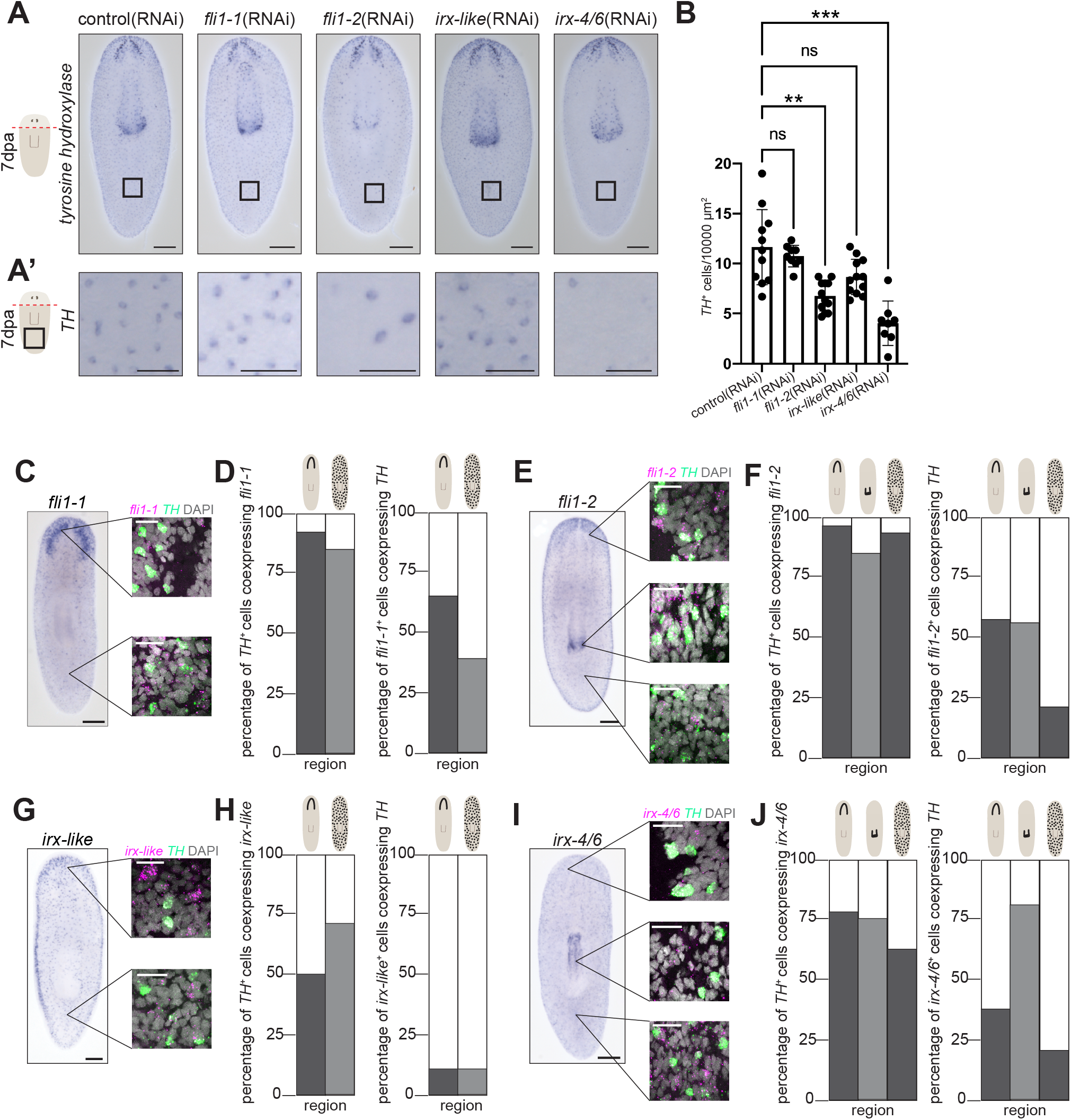
*FLI* and *IRX* genes are required for *TH^+^* cells in the PNS. A) Colorimetric ISH performed in our original screen showing *TH^+^*cells on the ventral side 7 days post head amputation after knockdown of *fli1-1, fli1-2, irx-like,* or *irx-4/6*. A’) Post-pharyngeal insets of A. B) Quantification of post-pharyngeal *TH^+^* cell number in three 10000 µm^2^ boxes. The average of each box is plotted on the graph. One-way ANOVA with Dunnett’s multiple comparisons test. C-J) Representative images and quantification of co-expression for *TH* with *fli1-1, fli1-2, irx- like,* and *irx-4/6*. Schematics above the bars indicate the region of the nervous system. For each gene, a colorimetric ISH is shown on the left, with insets showing co-expression of the gene of interest (magenta), *TH* (green), and DAPI (nuclei – gray) visualized after FISH. The degree of co-expression for each gene with *TH* is shown for the CNS and PNS for all genes and the PhNS for *fli1-2* and *irx-4/6*. (ns = not significant, ** P < 0.01, *** P < 0.001). Scale bars = 200 µm (A, C, E, G, I, except insets), 50 µm (A’), 20 µm (C, E, G, I insets).

First, we examined expression patterns of *FLI* and *IRX* genes. *fli1-1* is expressed in the brain, brain branches, and PNS^18^ and was co-expressed with *TH^+^ cells* in the CNS and PNS (Fig. 3C-D)*. fli1-2* is expressed broadly, with most *TH^+^* cells in all regions of the nervous system being *fli1-2^+^* (Fig. 3E-F). *irx-like* is expressed in a punctate pattern throughout the body, restricted to the ventral side, with co-expression noted in *TH^+^* cells of the CNS and PNS (Fig. 3G-H). *irx-4/6* is expressed in cells throughout the planarian body, including the pharynx. Most *TH^+^* cells also express *irx-4/6* (Fig. 3I-J). Our data indicate that *FLI* and *IRX* genes could impact *TH^+^* cells in a cell-autonomous manner, but that each gene is also likely to function in other cell types.

We next returned to investigating the function of FLI and IRX family members in regeneration and maintenance of dopaminergic neurons. Though we did not see a significant reduction of *TH^+^* cells in any location after perturbing *irx-like* or *fli1-1*, we hypothesized they could cooperate with *irx-4/6* and *fli1-2,* respectively. We completed double knockdowns of *IRX* or *FLI* pairs to see if there was an exacerbated phenotype. Indeed, we saw a stronger reduction of dopaminergic neurons in the tail in both double knockdown experiments (Fig. 4A-B, Extended Fig. 7A-D). Notably, we saw a bias in impacts to peripheral *TH^+^*cells across the dorsoventral axis. *irx-like*;*irx-4/6*(RNAi) caused a more severe reduction of dorsal peripheral dopaminergic neurons, while *fli1-1;fli1-2*(RNAi) depleted ventral peripheral dopaminergic neurons more strongly (Fig. 4A-B, Extended Fig. 7A-D). Therefore, we conclude that *irx-like, irx-4/6, fli1-1,* and *fli1-2* promote maintenance of peripheral dopaminergic neurons, with *IRX* playing a more penetrant role for dorsal *TH^+^* cells and *FLI* genes preferentially impacting ventral *TH^+^* cells.

**Figure 4:**
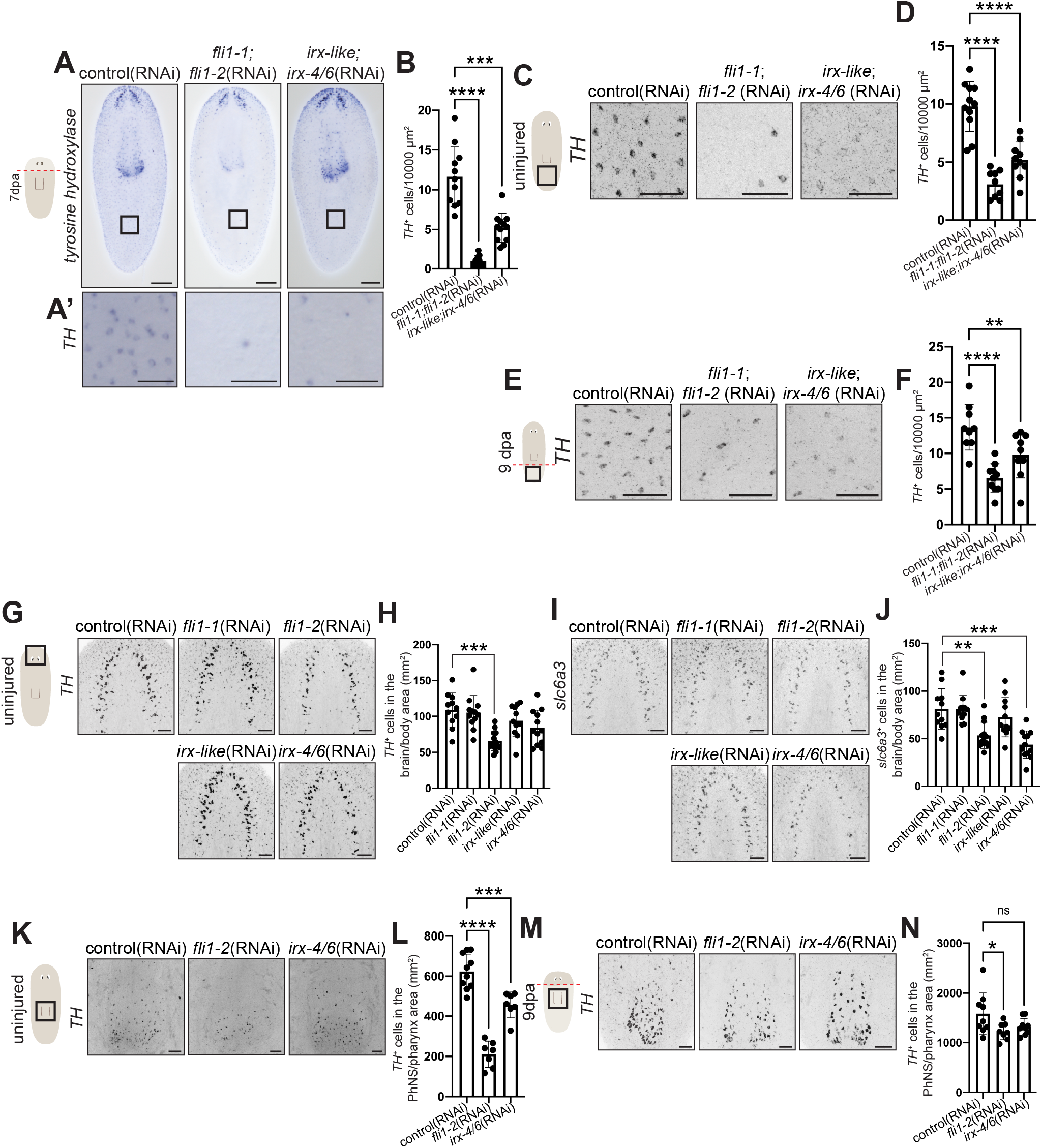
*fli1-2* and *irx-4/6* are required for *TH^+^* cells throughout the nervous systems. A) Whole-body colorimetric ISH focusing on the ventral side showing *TH^+^* cells in the maintained tail region 7 days post head amputation after double knockdown of *fli1-1;fli1-2* or *irx-like;irx- 4/6.* A’) Post-pharyngeal insets of A. B) Quantification of post-pharyngeal *TH^+^*cell number in three 10000 µm^2^ boxes. The average of each box is plotted on the graph. One-way ANOVA with Dunnett’s multiple comparisons test. C) Inverted grayscale FISH images showing *TH^+^* cells in the maintained tail after knockdown of *fli1-1;fli1-2* or *irx-like;irx-4/6* compared to control. D) Quantification of PNS *TH^+^* cell number in three 10000 µm^2^ boxes. The average of each box is plotted on the graph. One-way ANOVA with Dunnett’s multiple comparisons test. E) Inverted grayscale FISH images showing *TH^+^* cells in the regenerated tail 9 dpa after knockdown of *fli1- 1;fli1-2* or *irx-like;irx-4/6* compared to control. F) Quantification of PNS *TH^+^*cell number in three 10000 µm^2^ boxes. The average of each box is plotted on the graph. One-way ANOVA with Dunnett’s multiple comparisons test. G) Inverted grayscale FISH images showing *TH^+^* cells in the maintained tiara after knockdown of *fli1-1, fli1-2, irx-like,* or *irx-4/6* compared to control. H) Quantification of G. One-way ANOVA with Dunnett’s multiple comparisons test. I) Inverted grayscale FISH images showing *slc6a3^+^*cells in the maintained tiara after knockdown of *fli1-1, fli1-2, irx-like,* or *irx-4/6* compared to control. J) Quantification of I. One-way ANOVA with Dunnett’s multiple comparisons test. K) Inverted grayscale FISH images showing *TH^+^* cells in the maintained PhNS after knockdown of *fli1-2* or *irx-4/6* compared to control. L) Quantification of K. One-way ANOVA with Dunnett’s multiple comparisons test. M) Inverted grayscale FISH images showing *TH^+^*cells in the regenerated PhNS 9 dpa after knockdown of *fli1-2* or *irx-4/6* compared to control. N) Quantification of M. One-way ANOVA with Dunnett’s multiple comparisons test (ns = not significant, * P < 0.05, ** P < 0.01, *** P < 0.001, **** P < 0.0001). Scale bars = 200 µm (A except insets), 100 µm (K), 50 µm (A’, C, E, G, I, M).

We initially observed roles for *FLI* and *IRX* genes in *maintenance* of dopaminergic neurons in the non-regenerating posterior during head regeneration. We wanted to clarify whether these factors played roles in PNS maintenance or regeneration. Because we saw the strongest phenotype after double knockdowns, we investigated dopaminergic neuron maintenance after knockdown of *fli1-1;fli1-2* or *irx-like;irx-4/6* pairs. After RNAi, we examined changes in dopaminergic neurons using FISH to label *TH* and *slc6a3*. We detected a significant reduction of maintained peripheral *TH^+^*/*slc6a3^+^*cells in the absence of injury (Fig. 4C-D, Extended Fig. 7E-F). We then tested roles for *IRX* and *FLI* pairs in regeneration of peripheral neurons. After tail amputation, we observed that *fli1-1;fli1-2*(RNAi) and *irx-like;irx-4/6*(RNAi) animals had significantly fewer dopaminergic neurons in the regenerated PNS (Fig. 4E-F, Extended Fig.7G-H). We conclude that *fli1-1, fli1-2, irx-4/6,* and *irx-like* promote regeneration *and* maintenance of peripheral dopaminergic neurons.

The most obvious phenotypes we observed after perturbation *FLI* or *IRX* genes were in the PNS, but we noted expression of these genes in the brain and pharynx as well. We thus examined *TH* expression in the brain after knockdown of *fli1-1, fli1-2, irx-like,* or *irx-4/6.* Repeating the original paradigm with head amputation, we observed a consistent but non-significant decrease in *TH^+^/slc6a3^+^* cell number in the *regenerated* brain after knockdown of *FLI* or *IRX* genes, including double knockdown conditions (Extended Fig. 8A-H). However, we found that *fli1-2*(RNAi) and *irx-4/6*(RNAi) caused reductions in *TH^+^/slc6a3^+^* cells in the brain in the absence of injury (Fig. 4G-J). In this case, double knockdown of *fli1-1;fli1-2* or *irx-like;irx-4/6* did not exacerbate phenotypes (Extended Fig. 8I-L). We therefore concluded that *fli1-2* and *irx-4/6* are required for dopaminergic neuron *maintenance* in the brain, potentially through survival or preservation of the terminally differentiated fate.

Finally, *fli1-2* and *irx-4/6* are co-expressed with *TH* in the pharynx, so we sought to determine whether either gene played a role in pharyngeal dopaminergic neurons. After RNAi of *fli1-2* or *irx-4/6,* we completed FISH with *TH* and *laminin*^80^ to quantify pharynx area*. fli1-2*(RNAi) and *irx-4/6*(RNAi) led to significantly fewer dopaminergic neurons in non-regenerating pharynges relative to pharyngeal area (Fig. 4K-L, Extended Fig. 8M). We completed an alternate RNAi paradigm, this time observing the head as it regenerated the tail and pharynx. *fli1-2*(RNAi) caused a significant reduction of dopaminergic neurons in the regenerated PhNS after normalizing for pharynx size (Fig. 4M-N, Extended Fig. 8N).

Collectively, we determined that perturbating *FLI* or *IRX* family genes leads to reduction in dopaminergic cell number throughout the nervous system, in regeneration and maintenance. The effects we observed occur principally through the action of *fli1-2* and *irx-4/6*, but *fli1-1* and *irx-like* reinforced their paralogs in the PNS. Further, while *fli1-2* plays a role in *TH^+^* cells throughout the nervous system, *irx-4/6*’s role may be more specific to neuron maturation or terminal differentiation since *irx-4/6* is consistently required for maintenance of dopaminergic neurons. Together, we conclude that *fli1-2* and *irx-4/6* encode core transcription factors that drive dopaminergic neuron regeneration and maintenance throughout the planarian body.

### Functions for TH^+^ neurons in planarian movement

Dopaminergic neurons in planarians are responsible for muscle-mediated locomotion^39,81^. Given the identification of genes that impact dopaminergic neurons in different regions, we wanted to determine which *TH^+^* neurons are critical for animal movement. Planarians have a righting reflex, allowing them to flip over when placed ventral side up^82^. *TH*(RNAi) animals took significantly longer to flip over after regeneration (Fig. 5A). Perturbation of either *lmo1/3-1* or *app-L1,* which impact only CNS *TH^+^*cells, recapitulated this phenotype (Fig. 5A). In contrast, perturbations that predominantly impacted the PNS during head regeneration had milder impacts on the righting behavior. *fli1-2*(RNAi) caused a slight increase in time to flip over during regeneration and *irx4-6*(RNAi) did not change righting time (Fig. 5B). Our data suggest that *TH^+^* neurons in the brain are most important for muscle-mediated movement, while peripheral dopaminergic neurons may have additional unknown functions.

**Figure 5:**
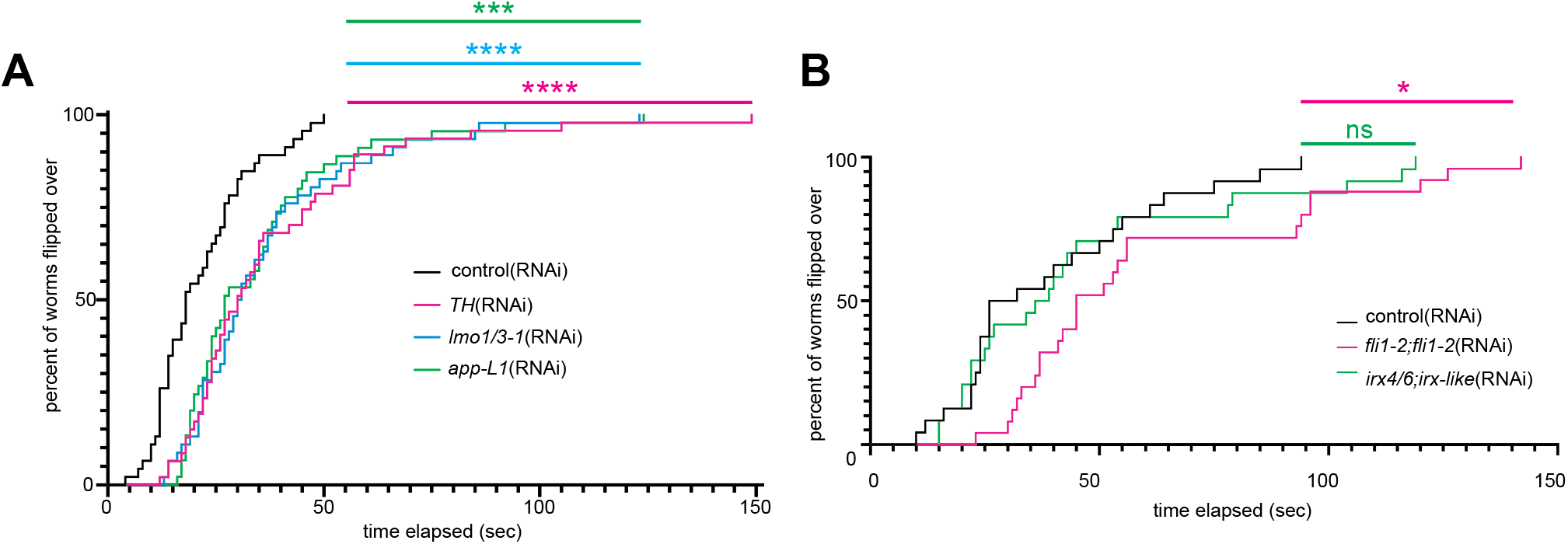
Ventral/dorsal flip is controlled by dopaminergic neurons in the CNS. A-B) Behavior of animals 6 dpa showing percentage of worms flipped over time across knockdown conditions. Analysis using Mantel-Cox log-rank test comparison between control and knockdown conditions. (ns = not significant, * P < 0.05, *** P < 0.001, **** P < 0.0001).

### soxB1-2 and foxA provide spatial direction for TH^+^ cells

Building on our identification of CNS-specific and body-wide dopaminergic neuron regulators, we next asked how peripheral and pharyngeal dopaminergic neurons are regionally specified. A hint came from our initial RNAi screen. Along with *fli1-2*(RNAi) and *irx4/6*(RNAi), we observed that *soxB1-2*(RNAi) impacted dopaminergic neurons in the PNS (Fig. 6A-B). Based on previous research*, soxB1-2* regulates sensory neurons and has been identified as an FSTF for neural progenitor populations^15,83^, but has not been implicated in regeneration of dopaminergic neurons, to our knowledge. We found that *soxB1-2*(RNAi) led to a significant reduction of peripheral *TH^+^/slc6a3^+^*cells in both regeneration and maintenance paradigms (Fig. 6C-F, Extended Fig. 9A-D). We further determined that *TH* is co-expressed with *soxB1-2* exclusively in the PNS, illustrating the location-specific influence of *soxB1-2* on dopaminergic neurons (Fig. 6G-H).

**Figure 6:**
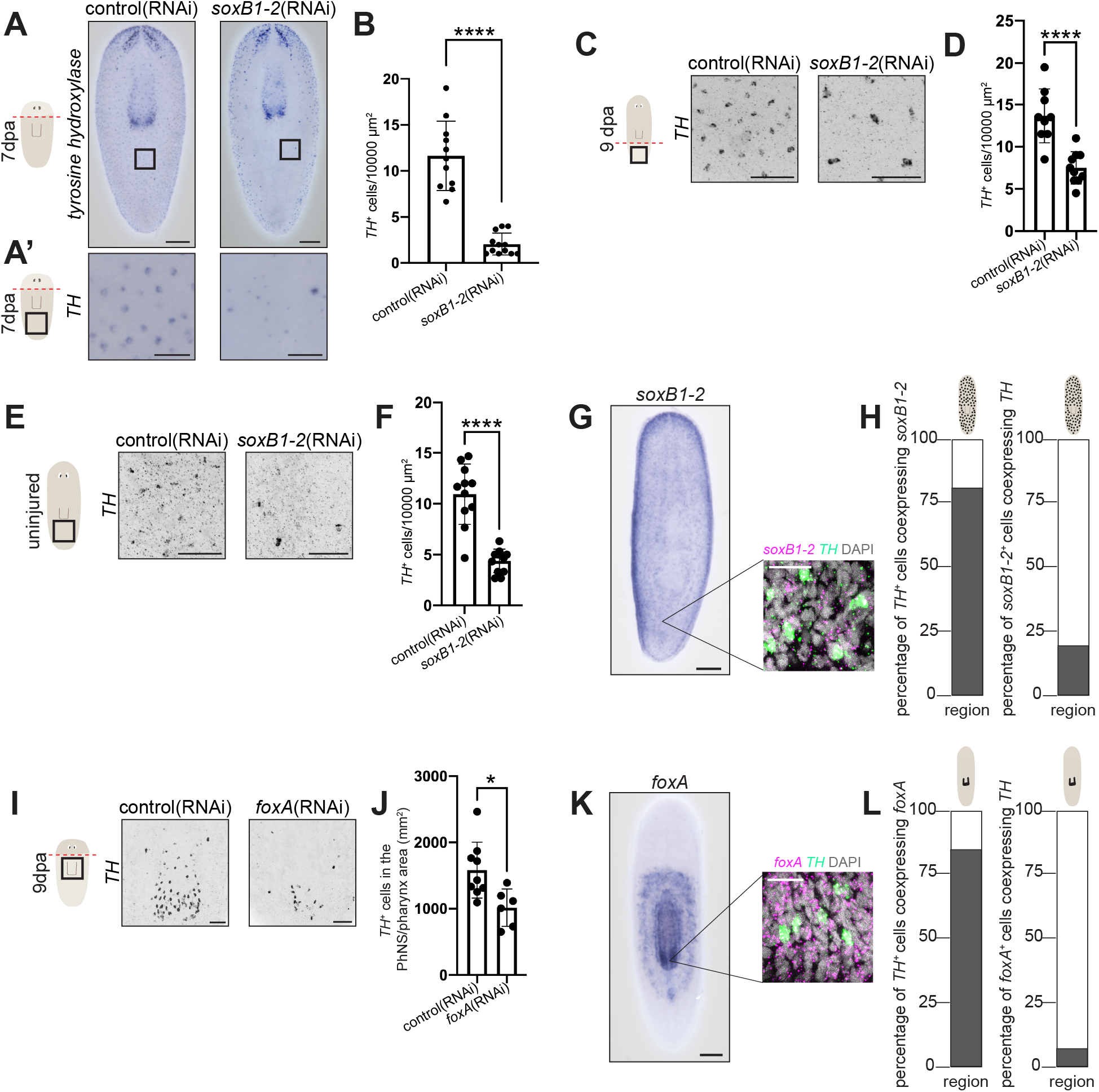
*soxB1-2* and *foxA* are location-specific regulators of *TH^+^* neurons. A) Colorimetric ISH performed in our original screen showing *TH^+^* cells on the ventral side 7 days post head amputation after knockdown of *soxB1-2* compared to control. A’) Post-pharyngeal insets of A. B) Quantification of post-pharyngeal *TH^+^* cell number in three 10000 µm^2^ boxes. The average of each box is plotted on the graph. Unpaired t-test. C) Inverted grayscale FISH images showing *TH^+^* cells in the regenerated tail 9 dpa after knockdown of *soxB1-2* compared to control. D) Quantification of PNS *TH^+^* cell number in three 10000 µm^2^ boxes. The average of each box is plotted on the graph. Unpaired t-test. E) Inverted grayscale FISH images showing *TH^+^* cells in the maintained tail after knockdown of *soxB1-2* compared to control. F) Quantification of PNS *TH^+^*cell number in three 10000 µm^2^ boxes. The average of each box is plotted on the graph. Unpaired t-test. G-H) Representative images and quantification of co-expression with *TH* and *soxB1-2.* Schematics above the bars indicate the region of the nervous system. A colorimetric ISH is shown on the left, with insets showing co-expression of *soxB1-2* (magenta), *TH* (green), and DAPI (nuclei – gray) visualized after FISH. The percentage of co-expression between *TH* and *soxB1-2* is shown for the PNS. I) Inverted grayscale FISH images showing *TH^+^* cells in the regenerated PhNS 9 dpa after knockdown of *foxA* compared to control. H) Quantification of G. Unpaired t-test. K-L) Representative images and quantification of co-expression for *TH* and *foxA.* Schematics above the bars indicate the region of the nervous system. A colorimetric ISH is shown on the left, with insets showing co-expression of *foxA* (magenta), *TH* (green), and DAPI (nuclei – gray) visualized after FISH. The percentage of cells co-expressing *TH* and *foxA* is shown for the PhNS. (* P < 0.05, **** P < 0.0001). Scale bars = 200 µm (A, G, K, except insets), 50 µm (A’, C, E, I), 20 µm (G, K insets).

When compiling our list of genes co-expressed with *TH* from single-cell experiments, we identified another region-specific transcription factor-encoding gene, *foxA*^14^. *foxA* is expressed pharyngeally and parapharyngeally and has been characterized for its unique role in pharyngeal regeneration^80,84^. We considered whether *foxA* might play a pharynx-specific role in promoting dopaminergic neurons. Indeed, we saw a significant reduction of *TH^+^* cells in regenerating pharynges after *foxA*(RNAi) (Fig. 6I-J, Extended Fig. 9E), even after normalizing to pharynx size, thus supporting a model in which *foxA* directs pharyngeal dopaminergic neuron fates. We were unable to study *foxA* effects in the homeostatic pharynx because *foxA*(RNAi) animals eject their pharynges. Supporting a role for *foxA* in pharyngeal *TH^+^* neurons, we observed PhNS-specific co-expression of *TH* with *foxA* (Fig. 6K-L). Collectively, our results indicate regenerative neurogenesis likely requires specific transcription factors that regulate production of neurons in the correct locations, with *soxB1-2* and *foxA* influencing PNS and PhNS dopaminergic neuron fates, respectively.

### Neuron identity and location are determined interdependently

Upon our discovery of a suite of key transcription factors that encode neuronal cell type *and* location for peripheral dopaminergic neurons, we were well positioned to test whether location and cell type are specified hierarchically or cooperatively. We first completed bulk RNA sequencing after RNAi targeting *soxB1-2, fli1-1;fli1-2,* or *irx-like;irx-4/6* (Fig. 7A). We identified hundreds of genes differentially expressed after *soxB1-2* knockdown (1682 transcripts downregulated and 820 transcripts upregulated compared to control), reflecting the broad function of this key transcription factor^83^ (Fig. 7B-C). We saw a much narrower subset of genes differentially expressed after *fli1-1;fli1-2*(RNAi) or *irx-like;irx4/6*(RNAi) (Fig. 7B-C), reflecting their narrower expression and more specific phenotypes. Notably, both double knockdowns led to more than 30 genes being differentially expressed in *TH^-^* cells, supporting roles of *FLI* and *IRX* genes as factors that influence dopaminergic neurons and other cell types^15,36^. Assessing all neural genes differentially expressed revealed a small but key set of overlapping genes, including *TH* and *slc6a3*, indicating a shared role of these genes in regulating dopaminergic neurons (Fig. 7D).

**Figure 7:**
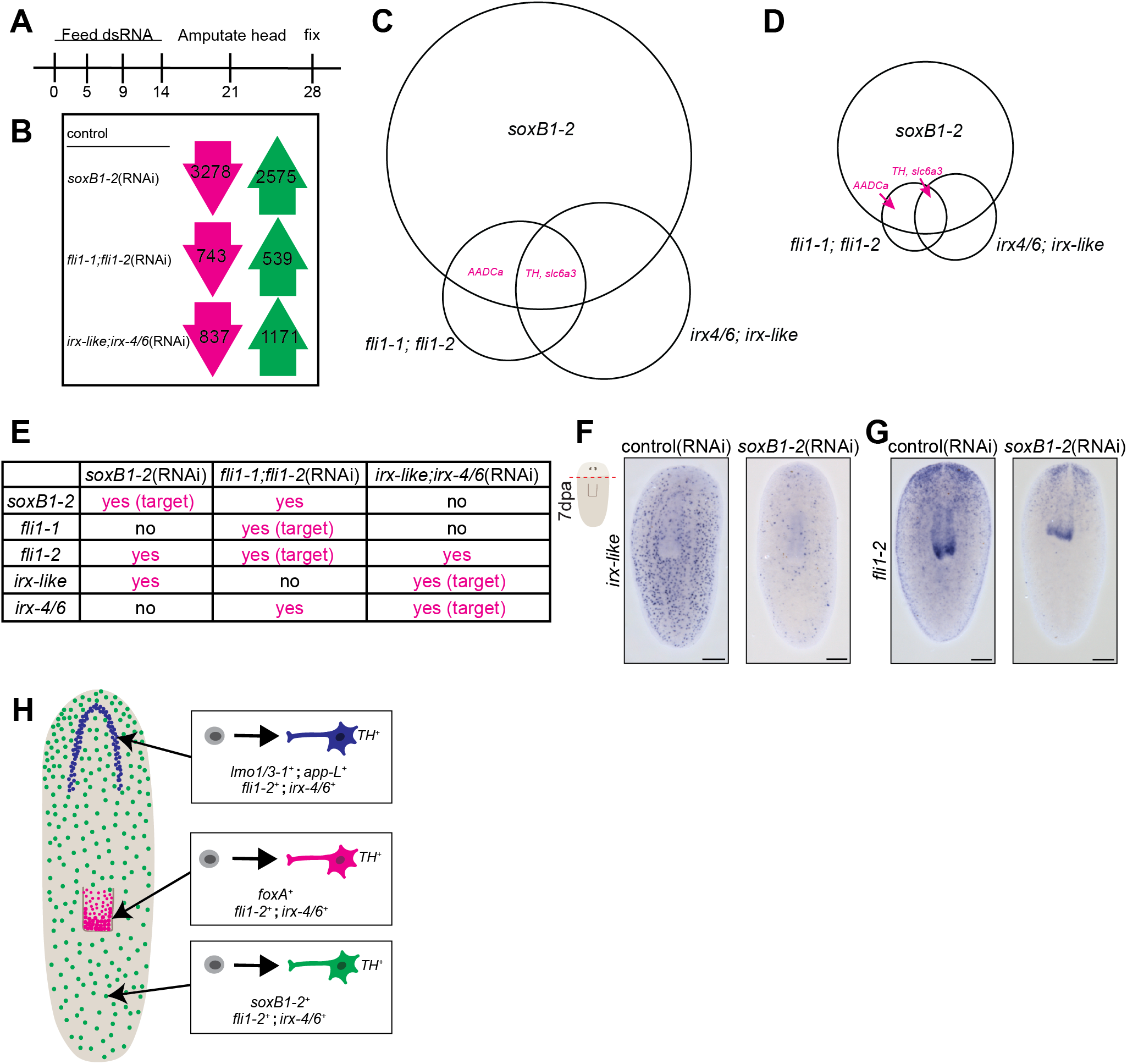
Identity and location of neurons are not determined in a hierarchical manner. A) RNAi paradigm for RNAseq experiments. B) Number of genes downregulated for each condition are shown in the magenta arrow (left) and genes upregulated for each condition in the green arrow (right). C) A Euler diagram depicting overlap for all genes significantly changed in each condition. D) A Euler diagram depicting overlap for all neural-enriched genes significantly changed in each condition. Neural genes were determined using single-cell transcriptomic data^14^. E) Table summarizing target genes from the RNAseq data, revealing whether the transcription factor-encoding genes influence transcript abundance for each other. “yes” (magenta) indicates the gene of interest (left) is significantly downregulated after the knockdown indicated (top). “no” indicates there is no significant change in the gene of interest’s expression. F-G) Whole-body colorimetric ISH showing expression of *irx-like* or *fli1-2* 7 days post head amputation after knockdown of *soxB1-2* compared to control. H) A schematic representation of our proposed model for the specification of *TH^+^*cell identity and location (CNS – blue, PNS – green, PhNS, magenta). Scale bars = 200 µm.

We next examined interdependence of *soxB1-2, fli1-1, fli1-2*, *irx-like*, and *irx4/6*. We determined through both ISH and RNA-sequencing that *soxB1-2*(RNAi) caused reduction in *irx- like* and *fli1-2* transcripts (Fig. 7E-G), which could support the idea that neural location is determined upstream of neurotransmitter identity. However, our data also revealed that double knockdown of *fli1-1;fli-2* resulted in a reduction of *soxB1-2* mRNA levels (Fig. 7E, Extended Fig. 10A). Further, double knockdown of *FLI* or *IRX* pairs resulted in a reduction of *irx-4/6* or *fli1-2*, respectively (Fig. 7E, Extended Fig. 10B-C). We conclude that there does not seem to be a definitive hierarchy in the creation of peripheral dopaminergic neurons. Instead, we argue that identity and location decisions can influence one another as they occur concurrently in each region of the planarian nervous system (Fig. 7H). Importantly, we note that regeneration-specific factors like *app-L1* may work to modify outcomes to influence cell fates after injury (Fig. 7H).

## Discussion

The ability to create new neurons in predictable types, numbers, and locations for a given injury or disease would revolutionize treatments of neurodegenerative diseases and other brain injuries. Our work aims to leverage highly regenerative animals to understand principles of successful neuron replacement. In this work, we focused on regeneration and maintenance of dopaminergic neurons in planarians. We discovered that two genes—*soxB1-2* and *foxA*—encode location of dopaminergic neurons within the PNS and PhNS respectively. In contrast, *fli1-2* and *irx-4/6* influence *TH^+^* cells throughout the nervous system. Based on previous research into neural progenitors, we confirm that *fli1-2* and *irx4/6* are expressed in G0 neural progenitors^15^ and that both *fli1-2* and *soxB1-2* are co-expressed with *smedwi-1*^36^. We thus conclude that these genes promote neurogenesis and maturation of progenitors into functional *TH*^+^ neurons. Further, the injury-specific role of *app-L1* indicates that other genes, including some that encode signaling molecules, can “tune” outcomes for regeneration, at least in the brain. Our results suggest that combinatorial programming of neurons from mammalian pluripotent cells might also require similar combination(s) of spatial- and identity-encoding factors to customize cell types for specific regions of the body and specific injuries.

Several key factors identified in this work have been shown to be important for dopaminergic neurons in other animals. *FLI* homologs specify dopaminergic neurons in *C. elegans*^85^. *foxA* homologs influence dopaminergic neuron birth, maintenance, and spontaneous age-related degeneration in murine models *in vitro* and *in vivo*^86–88^. The identification of conserved factors through our screen supports the idea that studying regeneration of cell types in planarians could reveal targets for directing mammalian neurogenesis *in vivo* or from pluripotent stem cells *in vitro*.

Importantly, we also identified genes *expressed* in dopaminergic neurons in other animals where a functional connection has not yet been established in the literature. According to single-cell sequencing data generated through human and mouse brains, mammalian homologs of *gst genes, caveolin genes, dzank1, lmo3*, and *app* are all expressed in dopaminergic neurons^68,89,90^. Further, *IRX* family genes are important for neurogenesis and neural differentiation in other organisms^91^ and have been implicated in Parkinson’s Disease^92^ but not dopaminergic neurons specifically. Our findings combined with results in the literature indicate that several genes of interest in our study could be interesting targets to explore in understanding dopaminergic neuron physiology and fate in other species. Importantly, our findings suggest that planarians are a suitable organism for discovery of novel genes impacting mammalian neurogenesis, neuron maturation, and neural maintenance.

Critically, we can now use our discoveries to reveal distinct mechanisms that support neuron formation, maturation, and survival. Future work will be needed to determine whether each gene is specific to regenerative neurogenesis and to determine whether they are required for cell survival. Understanding the cellular mechanisms involved in regulating dopaminergic neurons could aid in the application of our discoveries to other models. For example, we found that planarian *app-L1* plays a regeneration-specific role in *TH*^+^ neurons (Fig 2A-D, Extended Fig. 4D-F), potentially via neurogenesis. APP is best known for its pathological processing and aggregation in Alzheimer’s disease, with recent studies linking dysfunction of APP with degeneration of dopaminergic neurons in genetic models of Parkinson’s, potentially due to iron-induced toxicity^93–99^. One approach to dissecting *app-L1*’s role would be to use 6-OHDA^100–103^ to ablate dopaminergic neurons and dissect mechanisms leading to cell replacement versus large-scale tissue replacement. Previous research together with our study positions *app* as a primary candidate for future study as a regulator of neural regeneration, especially in the case of neurodegenerative diseases.

We also noted that *TH* is expressed in PNS neurons on both ventral and dorsal sides of the animal, and that *irx-like* was exclusively ventrally expressed. Planarians regulate dorsoventral identity through the action of *bone morphogenetic protein 4* (*bmp-4*) and *anti-dorsalizing morphogenetic protein* (*admp*)^104–108^. *bmp* is a dorsal regulator that inhibits *admp* expression while ventral *admp* activates *bmp* expression, creating a regulatory circuit^109^. Our work suggests that dorsal and ventral PNS neurons have distinct identities and functions. Identification of neuronal markers with polarized expression, like *irx-like*, can help elucidate distinct roles for regulation of populations of peripheral neurons. For example, distinct neurons could promote the beating of ventral cilia or the independent movement of dorsal and ventral muscles. Our work also provides a starting point from which to investigate polarity cues that provide location instructions for mature peripheral neurons. The spatial-regulating factors that we identified as critical for dopaminergic neurogenesis—*soxB1-2* and *foxA*— are likely regulated by well-known polarity cues that pattern the planarian body. Recent work identified roundabout receptor RoboA as a Slit-independent regulator of pharynx-specific genes, including *foxA*^84,110^. Further, the planarian CNS is positioned ventrally and the brain anteriorly; thus dorsoventral and anteroposterior cues could be upstream of pro-neural factors.

Altogether, we identified 11 genes that lead to targeted regeneration of dopaminergic neurons in distinct locations of the nervous system. The unprecedented level of molecular understanding of a single neuron type in planarians enabled us to begin defining the journey that pluripotent stem cells travel to adopt specific neuronal identity in distinct locations. We conclude that—in the absence of key developmental scaffolding—combinatorial mechanisms must concurrently specify neuronal identity and location during regeneration.

## Acknowledgements

We would like to thank previous members of the lab who assisted in cloning: Anna Iouchmanov, Justin McCurdy, and Raag Patel. Additionally, we would like to thank past and current Roberts-Galbraith lab members who assisted in project feedback and support over the years. We would like to thank Anthony Sego for his computational feedback and assistance in file conversions for behavioral data. We would also like to thank Dr. Muthugapatti Kandasamy and the Microscopy Core for training and technical support. We thank the Georgia Genomics and Bioinformatics Core at UGA for RNA-sequencing assistance. This work was initiated with support from the Alfred P. Sloan Foundation and the McKnight Foundation, through a Sloan Research Fellowship and a McKnight Scholars Award to RRG. Preliminary work on catecholamine neurotransmitters was initially supported by a CAREER award to RRG from the National Science Foundation under grant number 1942822. The study was broadened and completed through funding from the National Institute of Neurological Disorders and Stroke (NINDS) of the National Institutes of Health under award number 5R01NS128096. RRG is also supported by the Lois K. Miller Professorship at the University of Georgia (UGA). This research was supported in part by funds from the Lalita and Raghubir Sharma Distinguished Professorship in Toxicology to NMF. KBC was supported by the National Institute of General Medical Sciences of the National Institute of Health under award number 1T32GM142623 and NINDS under project number 5R25NS107179. KBC was supported by the University of Georgia Research Foundation and ARCS Foundation. RNG and OOO were both supported by funding from the Center for Undergraduate Research Opportunities and the Office of Experiential Learning at UGA. OOO was also supported by the Peach State Louis Stokes Alliance for Minority Participation funding through National Science Foundation grant 2110067. The content is solely the responsibility of the authors and does not necessarily represent the official views of funding agencies.

## Methods

### Animal Husbandry

Asexual planarians (*Schmidtea mediterranea*, CIW4 strain) were kept at 18°C in the dark in 1X Montjuïc salts^111^. Animals were housed in 9 cup Ziploc containers. They were fed beef liver paste (White Oak Pastures) every 1-2 weeks to maintain size and health. Prior to experimentation, animals were starved for a minimum of 7 days. Animals were periodically treated with gentamycin sulfate (Gemini Bio-Products) at concentration of 50 µg/mL to maintain health.

### Identification of genes of interest and cloning

We identified 74 genes of interest (Supplemental Table 1) enriched in *TH*^+^ neural subclusters 23, 26, and 37 from a single-cell sequencing atlas^14^. Most candidate genes were selected based on high log-fold enrichment, but we also selected a number of transcripts with lower enrichment that encoded transcription factors or signaling proteins. mRNA sequences were obtained from Planmine v3.0^112^. 20-22 bp primers were designed to amplify 500-750 bp of the transcript of interest using BatchPrimer3, Primer 3plus, or NCBI’s primer design tool^113–115^. We amplified genes of interest for cloning into EAM5112-digested pJC53.2 vector with standard molecular methods^22^. Sequence validation and orientation were determined after Sanger sequencing (GENEwiz/Azenta).

### Riboprobe Synthesis and *in situ* hybridization

Antisense riboprobes were synthesized using either SP6 or T7 polymerase using standard molecular methods^22^. All colorimetric *in situ* hybridization was completed using Digoxigenin-11-UTP (Roche)-labeled probes. Fluorescent *in situ* hybridization probes were synthesized using either Digoxigenin-11-UTP, Fluorescein-12-UTP (Roche), or DNP-11-UTP (Revvity Health Sciences). Colorimetric *in situ* hybridization (ISH) was performed as in King & Newmark, 2013^116^. Briefly, animals were killed using 7.5% N-Acetyl Cysteine in 1X PBS solution (1.37 M NaCl, 27 mM KCl, 100 mM Na_2_HPO_4_, 20 mM KH_2_PO_4_, pH 7.4) and fixed in 4% formaldehyde in PBSTx (PBS + 0.3% Triton-X100). They were then washed and dehydrated into methanol and stored at −20°C. Following rehydration, animals were washed, bleached (5% formamide and 1.25% H_2_O_2_ in 2.5% SSC. 1X SSC = 150 mM NaCl and 15 mM sodium citrate), and hybridized in riboprobe solution (riboprobe diluted in Hyb solution: 50% deionized formamide, 10% SSC, 1% Tween-20 in Diethyl Pyrocarbonate (DEPC)-treated water) with riboprobes diluted to between 1:800 and 1:4000 depending on the probe strength. After hybridizing for more than 16 hours, animals were washed and put into blocking solution (5% horse serum in TNTx: 0.1 M Tris pH 7.5, 0.15 M NaCl, 0.3% Triton-X-100) for one to two hours and then antibody solution (blocking solution with anti-digoxigenin Fab fragments conjugated with alkaline phosphatase, Roche). Samples were rocked at 4°C overnight. After eight TNTx washes, samples were developed using 5-Bromo-4-chloro-3-indolyl phosphate (BCIP, Roche) and nitro blue tetrazolium chloride (NBT, Roche) in alkaline phosphatase buffer (100 mM Tris pH 9.5, 100 mM NaCl, 50 mM MgCl_2_, 0.1% Tween-20, 10% PVA). Development was stopped with two five-minute PBSTx washes, followed by a twenty-minute wash in 100% ethanol. Ethanol was replaced with PBSTx and samples were cleared and mounted in 80% glycerol. Animals from ISH experiments examining *TH*^+^ PNS stains were mounted in PBSTx without clearing in 80% glycerol for ease of imaging dorsal or ventral cells. Some ISH experiments were conducted in an Insitu Pro Intavis hybridization robot.

For fluorescent ISH, we also used prior methods with minor changes^116^. Animals were processed using the colorimetric protocol up to the blocking step. Fluorescence blocking solution includes 0.5% Roche Western Blocking Reagent and 5% horse serum in TNTx, and the antibody solution includes probes synthesized with anti-DIG-POD (Roche) or anti-FITC-POD (Roche) at a dilution of 1:2000 or anti-DNP-HRP at a dilution of 1:1000. For double or triple fluorescent ISH, we used multiple probes with different labels (DIG, FITC, DNP). After washes, we complete one 10-minute development with TAMRA-tyramide [0.2% TAMRA-tyramide, 0.05% 4-IPBA, 0.05% H_2_O_2_ in TSA Buffer (2M NaCl in 0.1M Boric acid pH 8.5, filter sterilized)], animals were incubated in 0.1 M sodium azide in PBSTx for 45 minutes to inactivate the first antibody, followed by a second blocking step and incubation in the second antibody solution. Development with FAM-tyramide (0.2% FAM-tyramide, 0.05% 4-IPBA, 0.05% H_2_O_2_ in TSA Buffer) proceeds after 6 washes in TNTx. For triple fluorescent ISH, samples were again washed, incubated in sodium azide as above, then incubated in the third blocking step and incubation in the third antibody solution. Development with DyLight633 (0.3% DyLight633-tyramide, 0.05% 4-IPBA, 0.05% H2O2 in TSA Buffer) proceeded after 6 more washes in TNTx. Samples were stained overnight in DAPI solution (1ug DAPI/10mL TNTx) and then washed 6 times in TNTx. Fluorescent samples were mounted in Vectashield antifade mounting medium (Vector Labs).

For colorimetric images, slides were imaged on an Axiocam 506 color camera mounted on Zeiss Axio Zoom V.16 microscope using ZEN 2.3 pro software. We use a PlanNeoFluar Z 1x/0.25 FWD 56mm objective and magnification ranges from 32x to 63x for whole body images. For confocal imaging, a Zeiss LSM 880 confocal microscope was used with an upright AXIO imager Z2 and Zen Black 2.3 SPI software. Whole body images were captured using a 10x/0.3 objective and regional images were taken using the 20x/0.8 objective. For double fluorescent ISH, images were taken using 40x/1.4 or 63x/1.4 objectives and some samples were imaged using Zen 3.6 Blue software on a Zeiss LSM 900 Microscope with AI sample finder on an inverted Zeiss Axio Observer microscope stand. Images were processed using Photoshop and Illustrator (Adobe).

### Image Quantification

Data were either blinded manually by colleagues or using the automated plugin Blind Analysis Tools in Fiji (ZMBH Imaging Facility, University of Heidelberg). Brain-to-body ratios were quantified by tracing the cephalic ganglia and the body in triplicate in Fiji^18^. For dopaminergic CNS, bodies were measured as previously stated and the cells in the central tiara were counted using the Cell Counter ImageJ plugin^117^. Separately, the body area was measured through either tracing in triplicate or thresholding. Finally, a ratio of cell number to body area was calculated. For the PNS, cells per square area were counted. Image area and level of fluorescence were measured in FIJI (Fiji is Image J^118^). Graphing and statistical analysis were conducted using GraphPad Prism version 9.0.0 for Mac OS X, GraphPad Software, Boston, Massachusetts USA, www.graphpad.com (details in Figure legends). For dopaminergic CNS, bodies were measured as previously stated and the cells in the central tiara were counted using the Cell Counter ImageJ plugin^117^. Separately, the body area was measured through either tracing in triplicate or thresholding. Finally, a ratio of cell number to body area was calculated. For the PNS, cells per square area were counted. Image area and level of fluorescence were measured in FIJI (Fiji is Image J^118^). For the PhNS, cells in the pharynx were counted manually. The pharynx area was measured through tracing in triplicate. For quantifying co-expression, 10-20 cells were selected in each area of interest (CNS, PNS, PhNS) on a single channel throughout a z-stack. After cell selection, the channels were merged and cells expressing both channels were counted as co-expressing cells. At least four animals were used for each set of quantifications. Each data point graphed represents 40-80 cells. Graphing and statistical analysis were conducted using GraphPad Prism version 9.0.0 for Mac OS X, GraphPad Software, Boston, Massachusetts USA, www.graphpad.com (details in Figure legends).

### RNA interference (RNAi)

dsRNA of interest was synthesized from pJC53.2 clones using standard molecular techniques^119^. Animals were separated into 60x15 mm Petri dishes 2-3 days before feedings, with 10-12 worms 3-5 mm in length for each experiment. 5 μg of dsRNA was fed to worms along with 1 μl of McCormick green food dye and 30 μl beef liver paste. dsRNA matching *Aequorea victoria green fluorescent protein* (*GFP*) was used as the negative control and *smedwi-2* served as a positive control due to lethality of the phenotype^120^. For early experiments, *TH* was used as an additional positive control. Animals were allowed to eat for 2 hours at each feeding before they were washed and placed in a new petri dish with gentamycin antibiotic at a dilution of 1:1000 (50 mg/mL stock from Gemini Bio-Products). Further details for RNAi paradigms are included in figure legends. For double knockdown RNAi experiments, 5 μg of dsRNA for each gene of interest was combined for a total of 10 μg per feeding. All dsRNA was used at the same concentration for experimental and control conditions (500-1000 ng/μl). dsRNA was mixed with 30-50 μl of liver paste and 1 μl of green food dye for each feeding.

### Real Time Quantitative Polymerase Chain Reaction (RT-qPCR)

RNA was extracted from RNAi-treated animals using Trizol (Ambion) as per the manufacturer’s protocol. Purified RNA was reverse transcribed to cDNA using iScript cDNA synthesis kit (BioRad). When relevant, primers for RT-qPCR were designed to amplify a shorter segment (product size ∼100) of a transcript that does not overlap with the dsRNA construct (Supplemental Table 1). RT-qPCR was completed using three technical and biological replicates of each condition with a qPCR kit (Promega) using an Applied Biosystems QuantStudio3 Real-time qPCR machine. *β-tubulin* mRNA was used as a normalization transcript for all qPCR experiments^22^. RT-qPCR data were analyzed using GraphPad Prism 9. Statistical analyses were completed using unpaired t-tests (with Welch’s correction when standard error was not the same between samples).

### High performance liquid chromatography (HPLC)

RNAi-treated animals were weighed and then homogenized in 500ul 0.2N HClO_4_ on ice using a Branson Sonifer 250 with output control of 3, duty cycle 92%, and output 20 for 3 seconds before being stored on ice. Then, samples were centrifuged at 13,200 XG for 10 minutes at 4°C. An aliquot (20 µl) of the supernatant was injected into an HPLC (Waters Arc HPLC System™) with a Waters™ 3465 electrochemical detector (ECD) and the analytes were separated on a base-deactivated C18 column using an isocratic flow rate of 1 mL/min. The monoamine mobile phase was made as in previously reported^121^. Data dilution-adjusted data (1:25) were analyzed using GraphPad Prism 9.

### Behavior assay

For dorsal/ventral flip assay, animals were placed in a separate Petri dish in 1X Montjuïc salts on their dorsal side. Animals were observed under a microscope and timed until they “righted” themselves, returning to dorsal side up. If animals did not right themselves within five minutes, they were considered to have not flipped. Data were plotted as percent of worms flipped over in five second increments.

### Bulk RNA sequencing and analysis

We completed dsRNA synthesis, RNAi, and RNA purification as described above. *gfp* dsRNA was used as a negative control. For all knock down conditions (*gfp*, *fli1-1;fli1-2, irx-like;irx-4/6,* and *soxB1*-*2*), we completed five biological replicates with 10-11 animals per replicate. Libraries were constructed from submitted RNA samples using a KAPA Stranded RNA-Seq Kit, at a third volume of manufacturer’s instructions, and using XGen Normalase UDI primers to incorporate indexes into libraries. Resulting libraries were pooled equimolarly based on DNA concentrations determined through photospectropic measures of SYBR green and on average fragment size as determined by an Advanced Analytical Fragment Analyzer. The final pool was quantified with Qubit version 4 and using qPCR (KAPA Library Quant Kit), which was performed on a Roche LightCycler 480 II. Analysis of qPCR fluorescent data was performed in the LightCycler 480 software version 1.5.1.62 SP3. The distribution of DNA fragment sizes of the final pool was quantified with an Advanced Analytical Fragment Analyzer. The pooled libraries were sequenced on an Illumina NextSeq 2000 using a P3_SBS 200 cycle reagent kit. Adapter trimming of reads was performed in Illumina BaseSpace.

Reads were mapped using CLC Genomics Workbench 24.0.1. All runs were mapped to the Dresden (dd_Smed_v6) transcriptome^14^ using default settings and Batch mode. Paired distances were auto-detected, with a max number of hits at 10, and “reverse” for strand setting. Expression values were reported as RPKM (reads per kilobase of transcript per million mapped reads). ≥80% of paired reads were mapped to this reference per run. After mapping, we performed differential gene expression for RNA-Sequencing. We tested differential expression due to condition, comparing each experimental RNAi condition to the control group *gfp*(RNAi). Raw reads will be available on NCBI Sequence Read Archive. Differential expression data is available in Supplemental Table 2.

For comparing differentially expressed genes (P-value of < 0.05) in each knockdown condition, Euler diagrams were assembled in R Studio 2024.04.01 using the eulerr package^122^. We also determined which differentially expressed genes were neural-enriched using Neural+Brain data from Fincher et al., 2018^14^.

### Protein alignment and phylogenetic analysis

Alignment and analysis were completed as previously described^71,123^. Briefly, FASTA sequences for planarian transcripts were obtained from available transcriptomes in PlanMine^112^ and translated to the longest open reading frame with Expasy^124^. Reference sequences for putative homologs in other organisms were obtained from NCBI Gene database^125^. Planarian protein sequences were compared to reference sequences in a phylogenetic analysis using phylogeny.fr “one-click mode” on default settings with MUSCLE alignment and G-blocks curation^126^. Additionally, for naming conventions, planarian protein sequences were loaded into phylogeny.fr “BLAST explorer” to create a phylogenetic tree of similar proteins from the “non redundant protein” database in NCBI from 2021-02-27 with an e value cut off of 1.e-5^127^. The top homolog in mice and human was also BLAST searched in PlanMine to confirm nomenclature. Relevant protein domains were confirmed by analyzing the longest open reading frame using InterProScan^128^. See supplemental figure 1-3 for more information.

## Extended figure legends

**Extended Figure 1:**
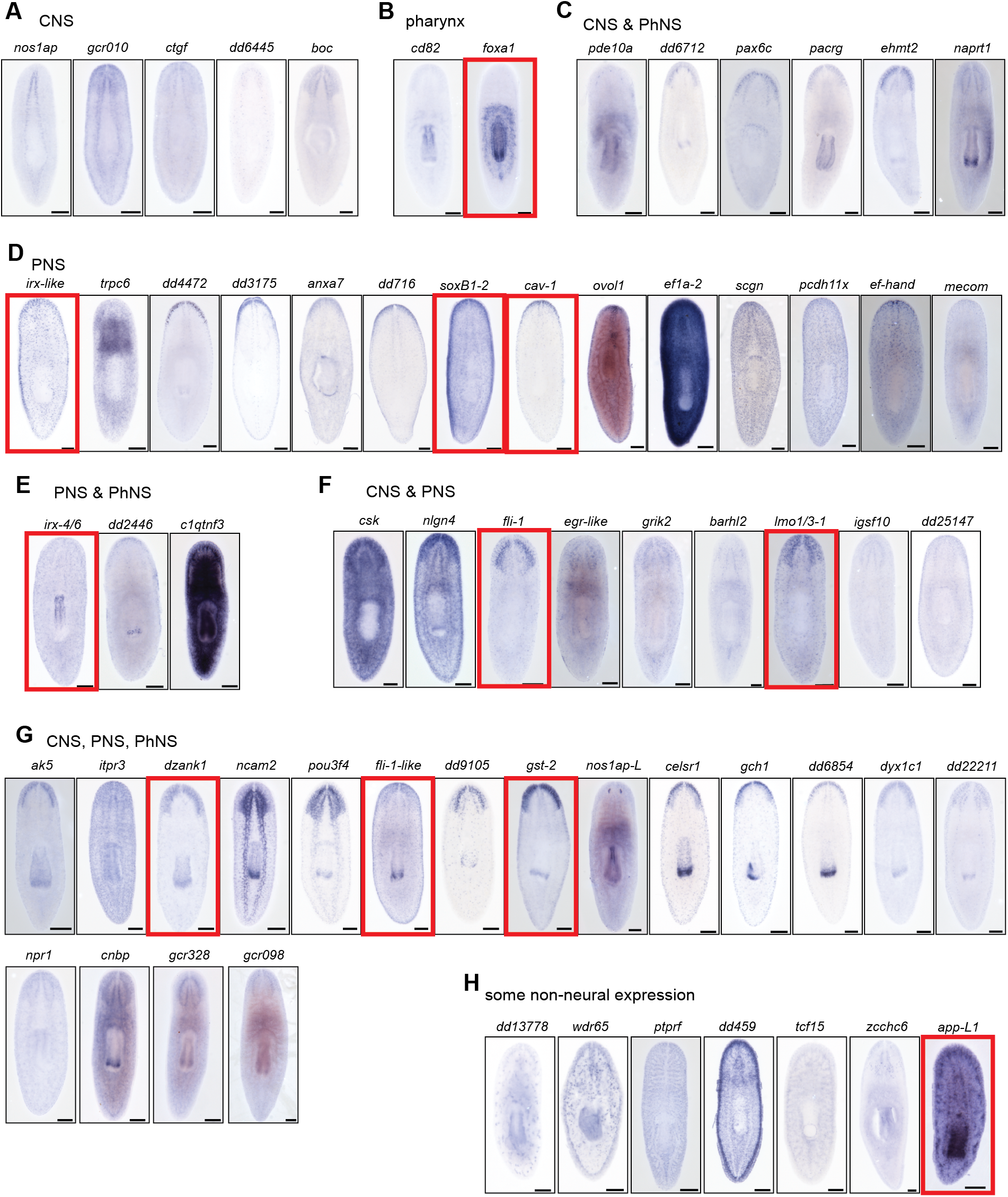
Expression patterns of genes from the screen. A-G) Genes expressed in different regions of the nervous system. H) Genes with both neural and obvious non-neural expression. Red boxes denote genes for which RNAi impacted *TH*^+^ cells (See Fig. 1N). Scale bars = 200 µm.

**Extended Figure 2:**
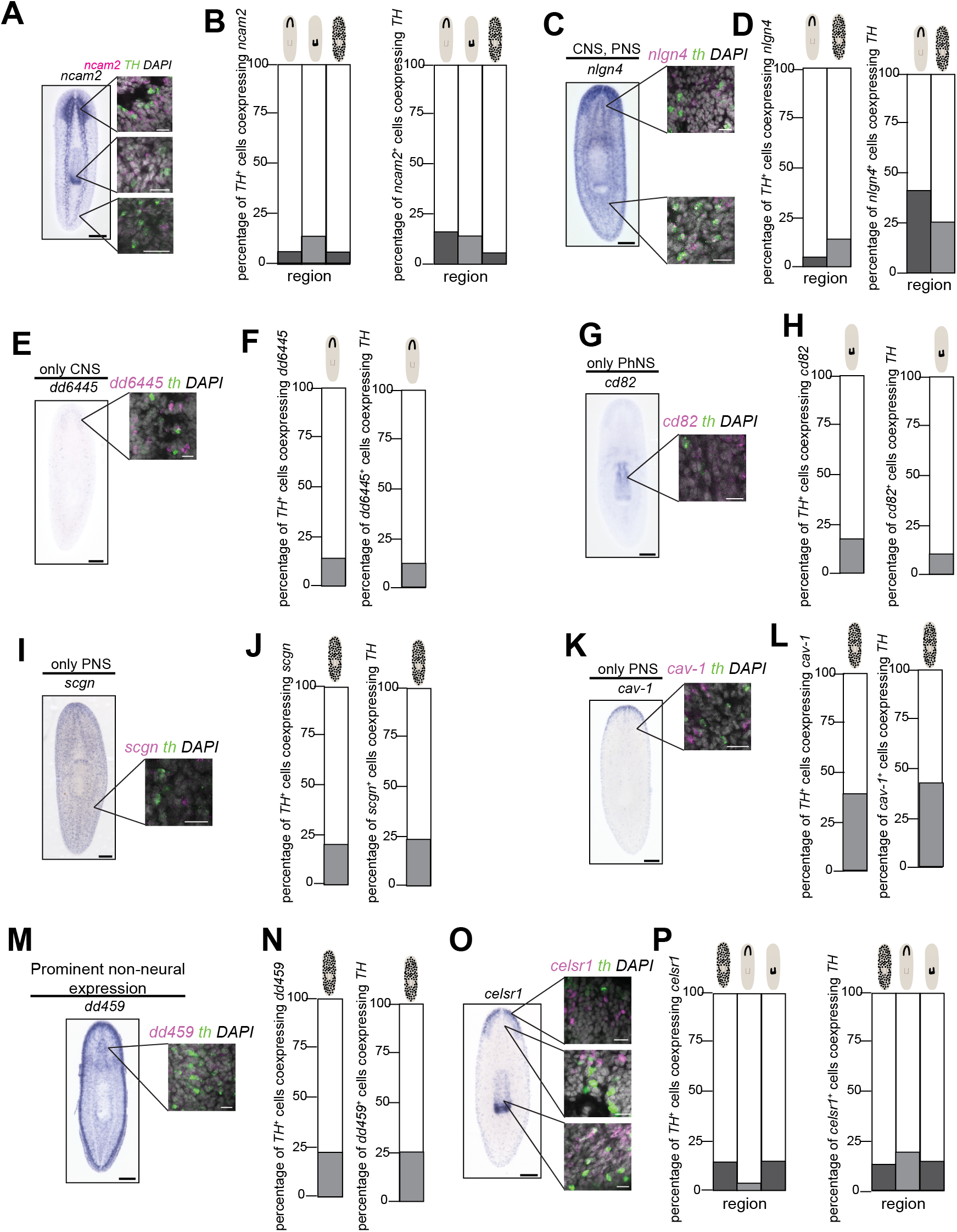
Additional co-expression of *TH* with genes from the screen. A-O) Representative images and quantification of co-expression with *TH* for eight genes (*ncam2, nlgn4, dd6445, cd82, scgn, cav-1, dd459,* and *celsr1*) found in different regions of the nervous system. Schematics above the bars indicate the region of the nervous system. For each gene, a colorimetric ISH is shown on the left, with insets showing co-expression of the gene of interest (magenta) and *TH* (green) visualized after FISH. Quantification of co-expression for each gene and *TH*. Scale bars = 200 µm (colorimetric), 20 µm (insets).

**Extended Figure 3:**
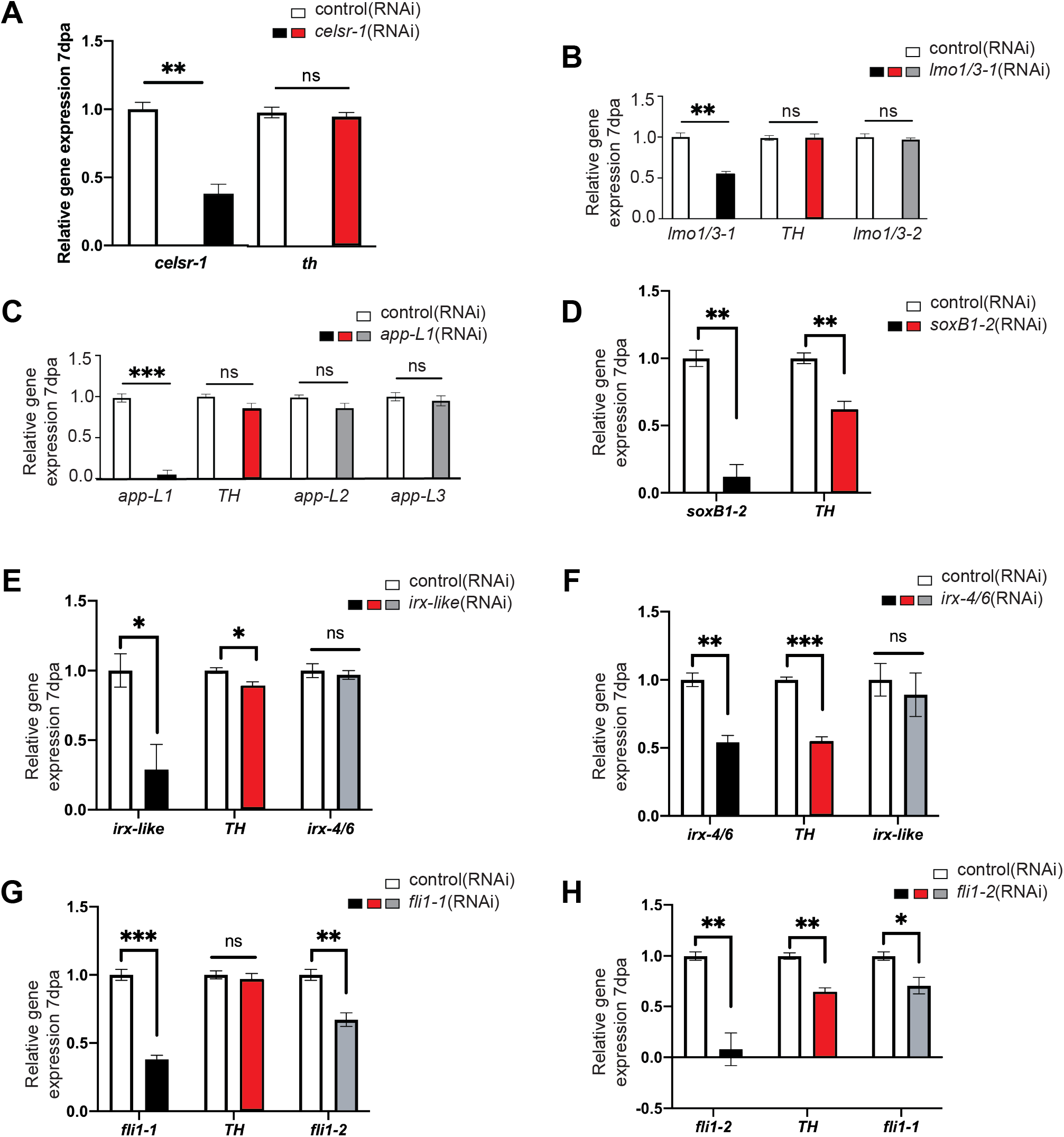
RT-qPCR confirming knockdown efficiency, target specificity, and level of *TH* in a few select genes. A-H) Relative gene expression of genes of interest (x-axis) 7 dpa after knockdown of selected candidate genes. For each RNAi, there were at least 3 biological replicates. RT-qPCR was completed using biological and technical triplicates. An unpaired t-test was used for all statistical analyses (ns = not significant, * P < 0.05, ** P < 0.01, *** P < 0.001). Black bar graphs = knockdown checks for indicated candidate genes. Red bar graphs = evaluation of *TH* transcript. Gray bar graphs = specificity checks with paralogous genes.

**Extended Figure 4:**
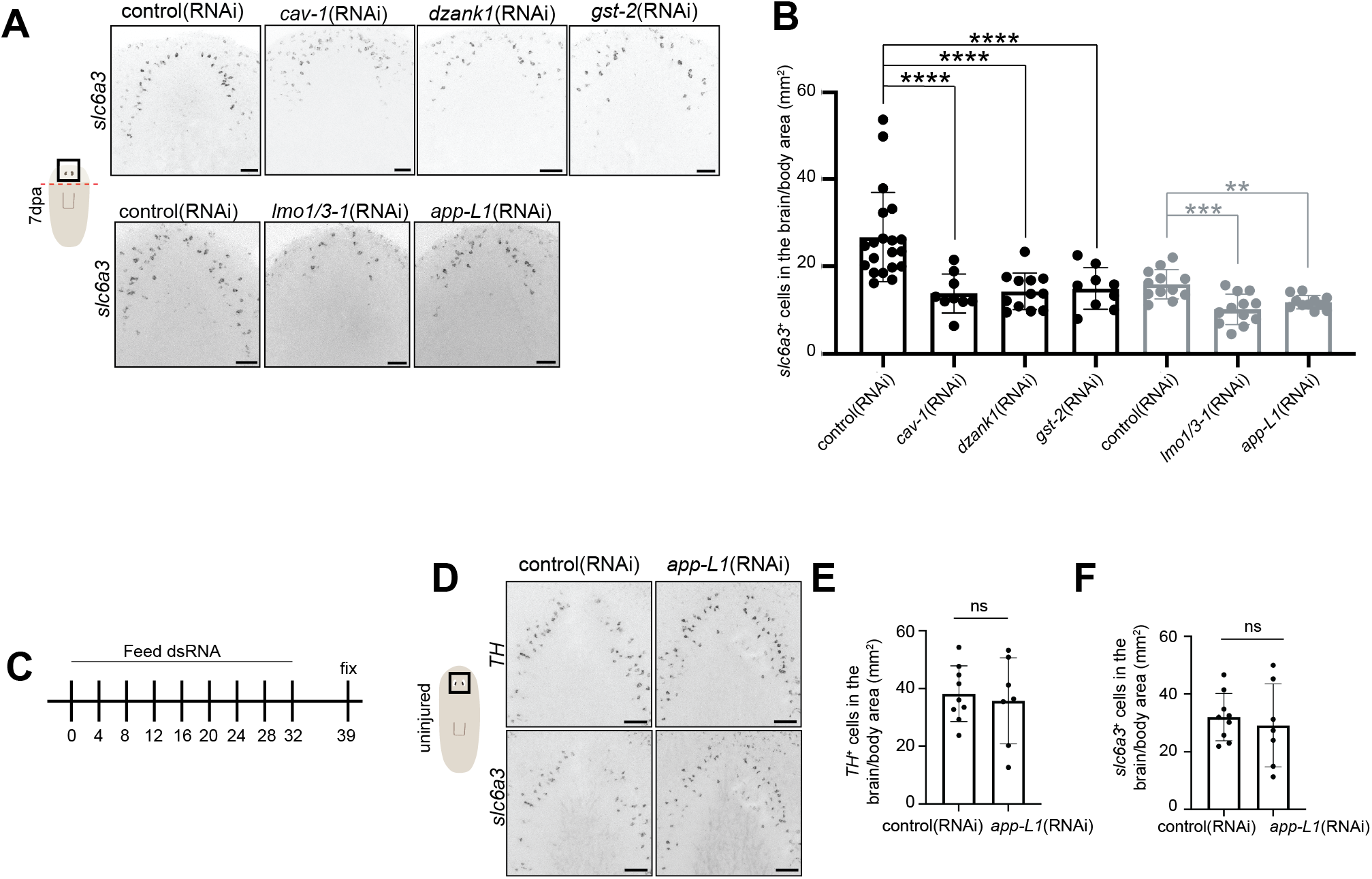
*lmo1/3-1* and *app-L1* are location-specific. A) Inverted grayscale FISH images showing *slc6a3*^+^ cells in the regenerated tiara 7 dpa after knockdown of *cav-1, dzank1, gst-2, lmo1/3-1* or *app-L1*. B) Quantification of animals from A. Left - One-way ANOVA with Dunnett’s multiple comparisons test, right – Brown-Forsyth and Welch ANOVA with Dunnett’s T3 multiple comparisons test. C) Long-term maintenance RNAi paradigm. D) Inverted grayscale FISH images showing either *TH*^+^ (top) or *slc6a3*^+^ cells (bottom) in the maintained tiara after knockdown of *app-L1*. E-F) Quantification of D. Unpaired t-test.

**Extended Figure 5:**
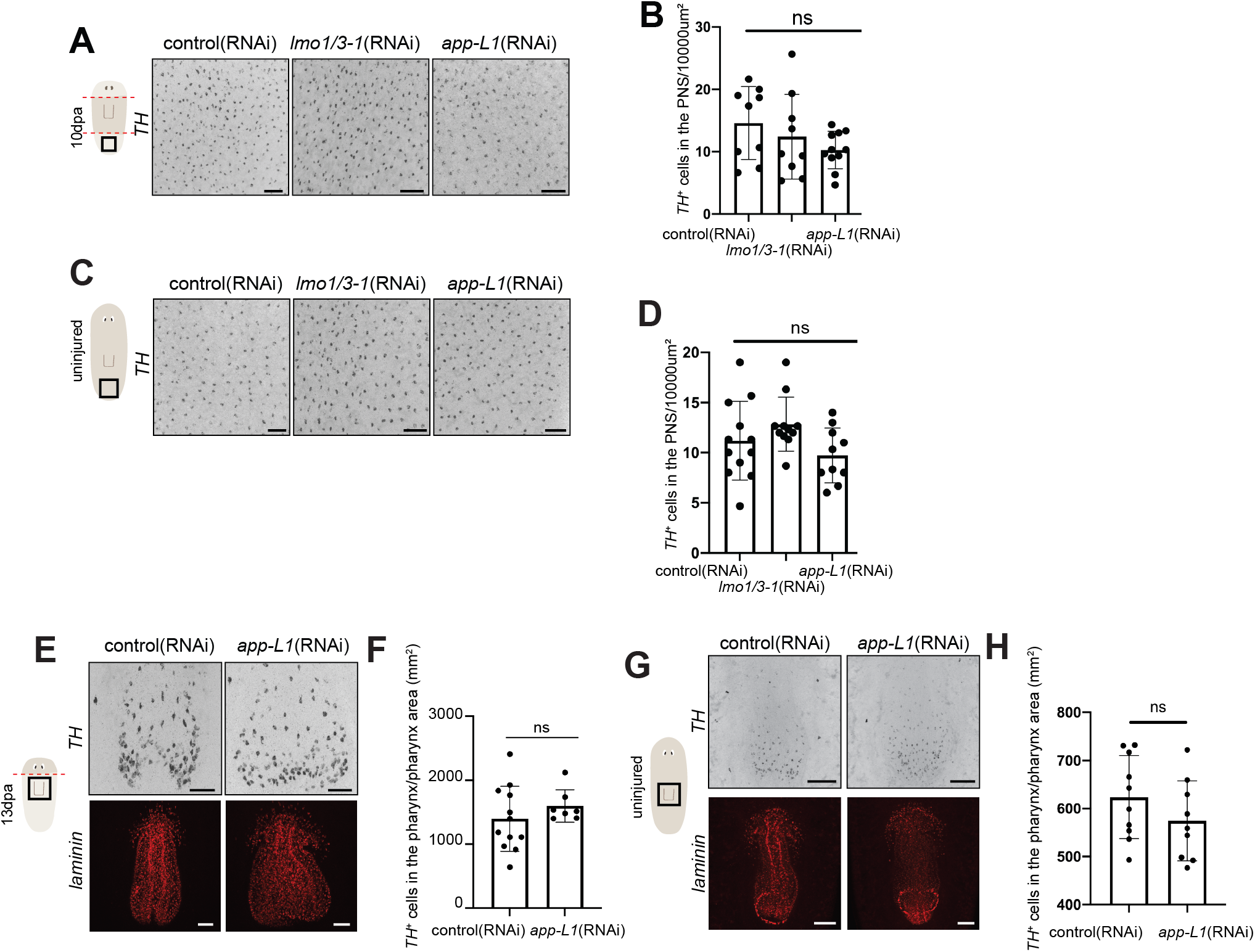
*lmo1/3-1* and *app-L1* do not affect the PNS or PhNS. A, C) Inverted grayscale FISH images showing either *TH*^+^ cell number in regenerated tails 10 days post trunk amputation (A) or uninjured tails (C) after knockdown of *lmo1/3-1* or *app-L1* compared to control. B, D) Quantification of A and C, respectively. One-way ANOVA with Dunnett’s multiple comparisons test (ns = not significant). E) FISH images showing *TH*^+^ cells (top, inverted grayscale) in regenerated pharynges (marked with *laminin*, bottom) 13 dpa after knockdown of *app-L1* compared to control. F) Quantification of *TH*^+^ cells in E normalized to pharynx size. Unpaired t-test. G) FISH images showing *TH*^+^ cells (top, inverted grayscale) in maintained pharynges (marked with *laminin*, bottom) after knockdown of *app-L1* compared to control. H) Quantification of *TH*^+^ cells in G normalized to pharynx size. Unpaired t-test. (ns = not significant) Scale bars = 100 µm (G) 50 µm (A, C, E).

**Extended Figure 6:**
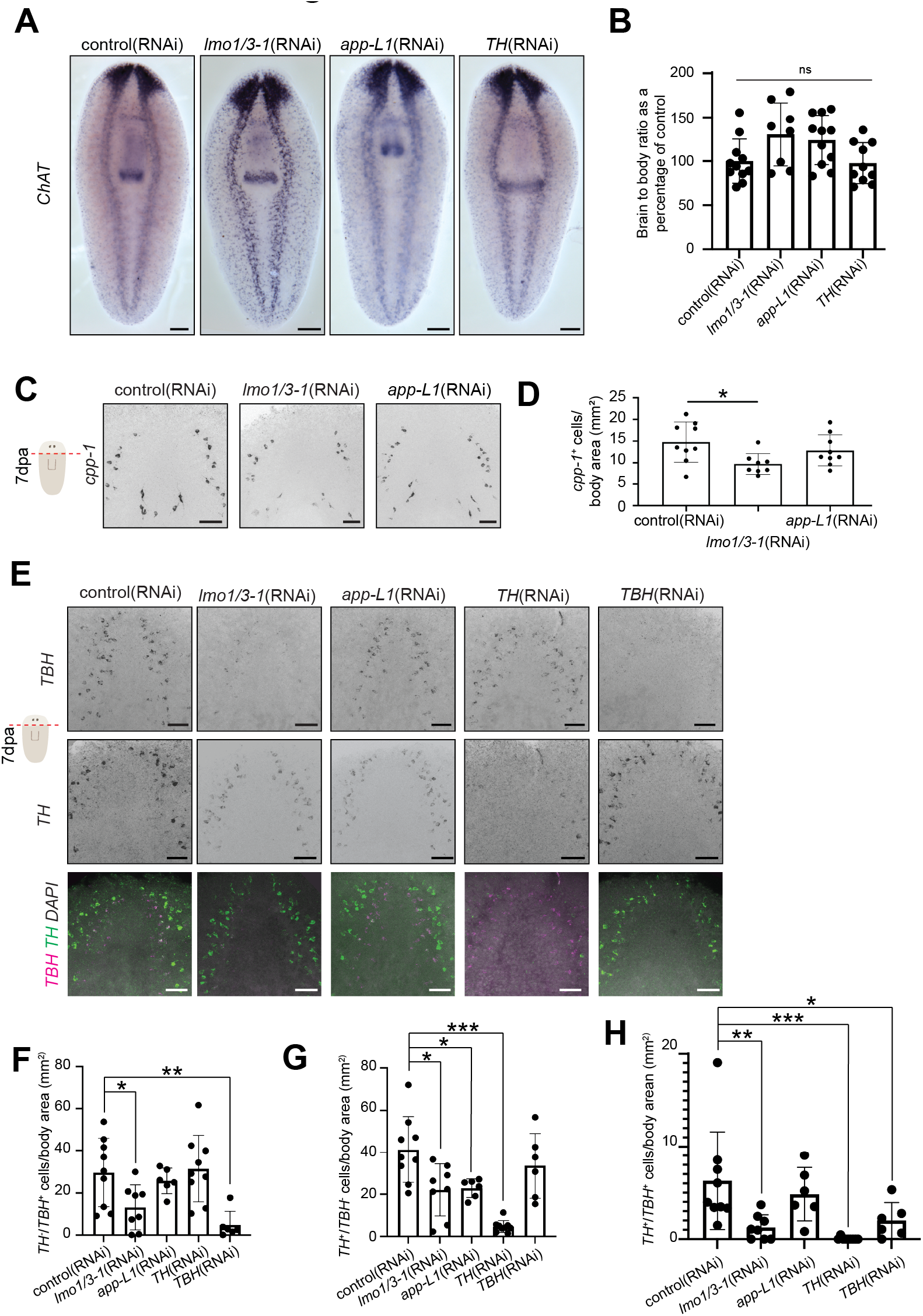
Specificity of *lmo1/3-1* and *app-L1*. A) Colorimetric ISH showing regenerated brain size (marked with *ChAT*^79^) 7 dpa after knockdown of either *lmo1/3-1* or *app-L1* or *TH* compared to control. B) Quantification of A. One-way ANOVA and Dunnett’s multiple comparisons test. C) Inverted grayscale FISH images showing *cpp-1*^+^ cells in 7 dpa regenerated heads after knockdown of *lmo1/3-1* or *app-L1* compared to control. D) Quantification of C. One-way ANOVA and Dunnett’s multiple comparisons test. E) FISH images showing either *TBH* (top – inverted grayscale), *TH* (middle – inverted grayscale), or both (bottom - *TBH* magenta, *TH* green, DAPI gray) in 7 dpa regenerated heads after knockdown of *lmo1/3-1*, *app-L1, TH*, or *TBH* compared to control. F-H) Quantification of E. F) Brown-Forsyth and Welch ANOVA with Dunnett’s T3 multiple comparisons. G-H) One-way ANOVA with Dunnett’s multiple comparisons test. (ns = not significant, * P < 0.05, ** P < 0.01, *** P < 0.001). Scale bars = 200 µm (A), 50 µm (C, E).

**Extended Figure 7:**
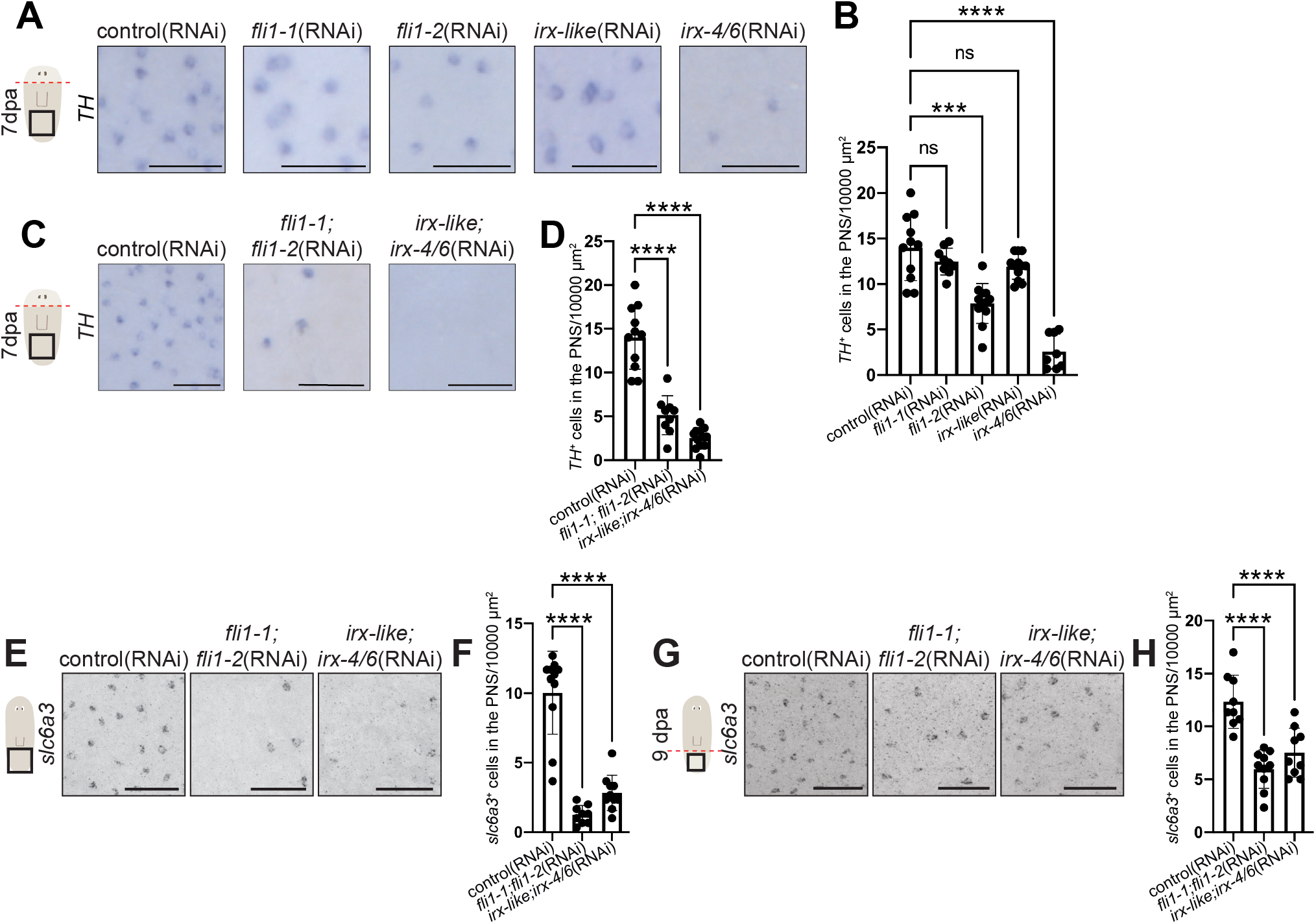
*FLI* and *IRX* genes are required for regeneration and maintenance of PNS dopaminergic neurons. A) Post-pharyngeal insets of colorimetric ISH imaged on the dorsal side showing *TH^+^* cells in the maintained tail 7 days post head amputation after knockdown of *fli1-1, fli1-2, irx-like* or *irx- 4/6* compared to control. B) Quantification of PNS *TH^+^* cell number in three 10000 µm^2^ boxes. The average of each box is plotted on the graph. One-way ANOVA with Dunnett’s multiple comparisons test. C) Post-pharyngeal insets of colorimetric ISH imaged on the dorsal side showing *TH^+^* cells in the maintained tail 7 days post head amputation after double knockdown of *fli1-1;fli1-2* or *irx-like;irx-4/6* compared to control. D) Quantification of PNS *TH^+^* cell number in three 10000 µm^2^ boxes. The average of each box is plotted on the graph. One-way ANOVA with Dunnett’s multiple comparisons test. E) Inverted grayscale FISH images showing *slc6a3^+^*cells in the maintained tail after double knockdown of *fli1-1;fli1-2* or *irx-like;irx-4/6* compared to control. F) Quantification of PNS *slc6a3^+^*cell number in three 10000 µm^2^ boxes. The average of each box is plotted on the graph. One-way ANOVA with Dunnett’s multiple comparisons test. G) Inverted grayscale FISH images showing *slc6a3^+^* cells in the regenerated tail 7 dpa after double knockdown of *fli1-1;fli1-2* or *irx-like;irx-4/6* compared to control. H) Quantification of PNS *slc6a3^+^* cell number in three 10000 µm^2^ boxes. The average of each box is plotted on the graph. One-way ANOVA with Dunnett’s multiple comparisons test. (ns = not significant, *** P < 0.001, **** P < 0.0001). Scale bars = 200 µm (A, C), 50 µm (E, G)

**Extended Figure 8:**
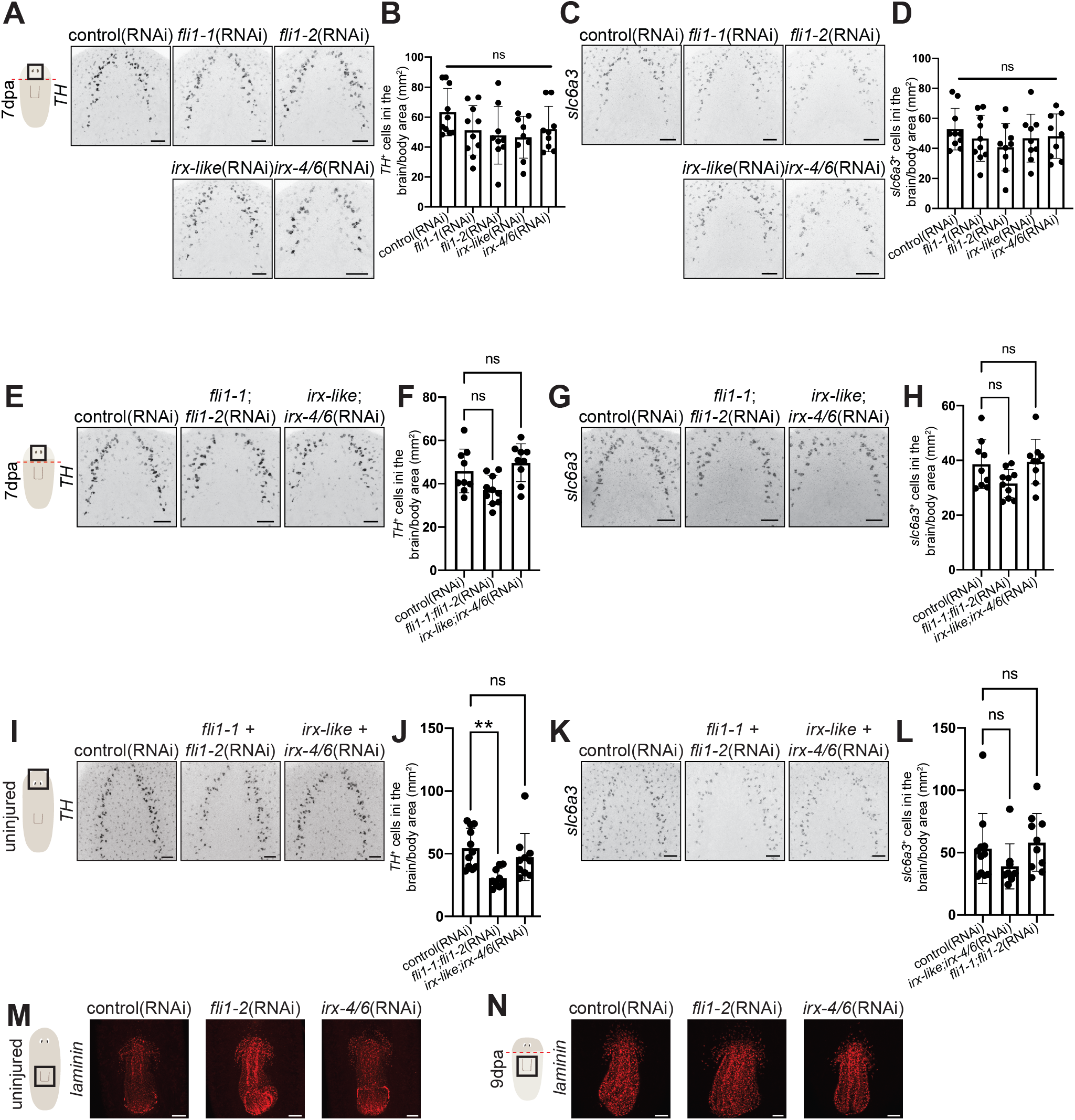
*FLI* and *IRX* genes regulate *TH^+^* cells in the CNS and PhNS. A) Inverted grayscale FISH images showing *TH^+^* cells in the regenerated brain 7 dpa after knockdown of *fli1-1, fli1-2, irx-like,* or *irx-4/6* compared to control. B) Quantification of A. One-way ANOVA with Dunnett’s multiple comparisons test. C) Inverted grayscale FISH images showing *slc6a3^+^* cells in the regenerated brain 7 dpa after knockdown of *fli1-1, fli1-2, irx-like,* or *irx-4/6* compared to control. D) Quantification of C. One-way ANOVA with Dunnett’s multiple comparisons test. E) Inverted grayscale FISH images showing *TH^+^* cells in the regenerated brain 7 dpa after double knockdown of *fli1-1;fli1-2* or *irx-like;irx-4/6* compared to control. F) Quantification of E. One-way ANOVA with Dunnett’s multiple comparisons test. G) Inverted grayscale FISH images showing *slc6a3^+^*cells in the regenerated brain 7 dpa after double knockdown of *fli1-1;fli1-2* or *irx-like;irx-4/6* compared to control. H) Quantification of G. One-way ANOVA with Dunnett’s multiple comparisons test. I) Inverted grayscale FISH images showing *TH^+^* cells in the maintained brain after double knockdown of *fli1-1;fli1-2* or *irx-like;irx- 4/6* compared to control. J) Quantification of I. One-way ANOVA with Dunnett’s multiple comparisons test. K) Inverted grayscale FISH images showing *slc6a3^+^*cells in the maintained brain after double knockdown of *fli1-1;fli1-2* or *irx-like;irx-4/6* compared to control. L) Quantification of K. One-way ANOVA with Dunnett’s multiple comparisons test. M) FISH images showing *laminin* expression in maintained pharynges after knockdown of *fli1-2* or *irx-4/6* compared to control. N) FISH images showing *laminin* expression in regenerated pharynges 9 dpa after knockdown of *fli1-2* or *irx-4/6* compared to control. (ns = not significant, ** P < 0.01). Scale bars = 100 µm (N), 50 µm (A, C, E, G, I, K, M)

**Extended Figure 9:**
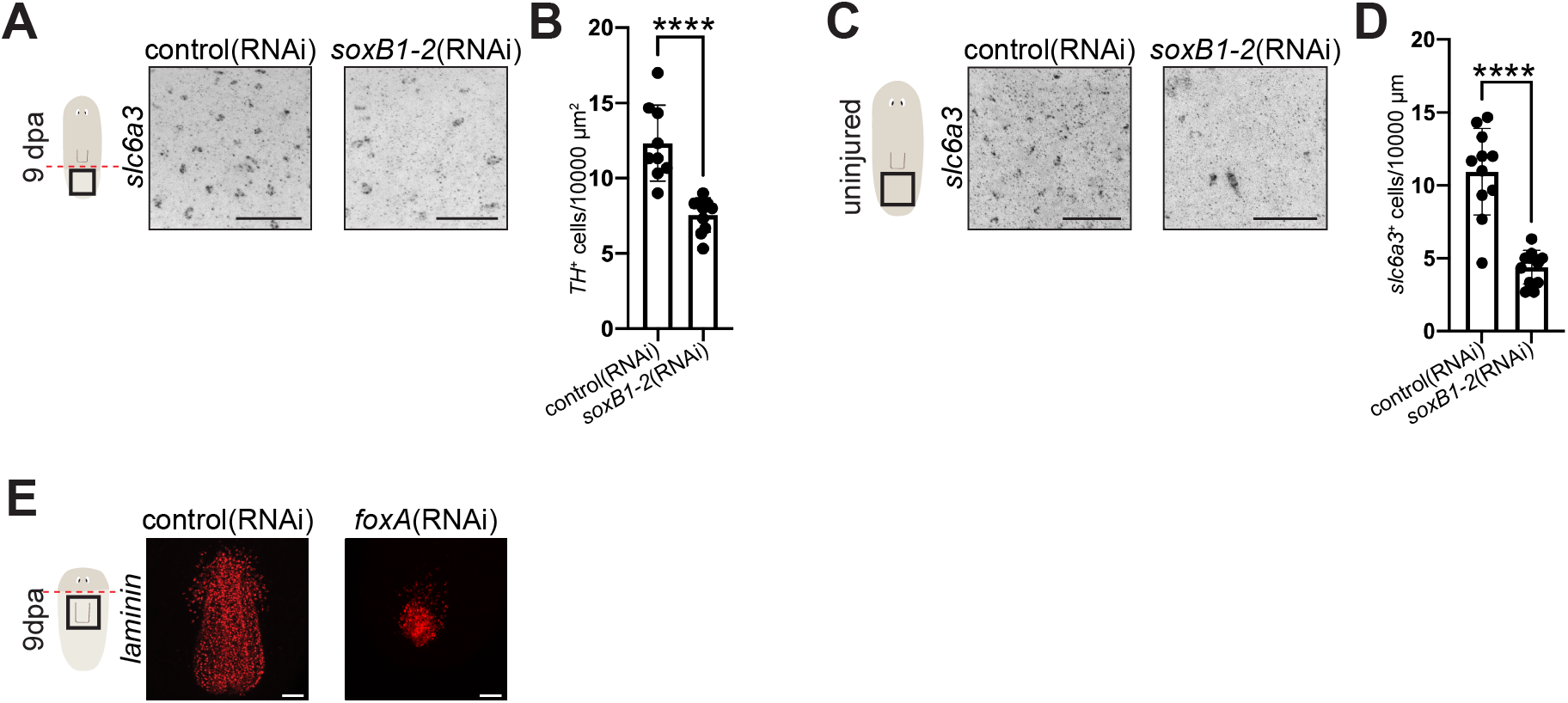
*soxB1-2* and *foxA* regulate dopaminergic neurons in a location-specific manner. A) Inverted grayscale FISH images showing *slc6a3^+^*cells in regenerated tails 9 dpa after knockdown of *soxB1-2* compared to control. B) Quantification of cell number in three 10000 µm^2^ boxes. The average of each box is plotted on the graph. Unpaired t-test. C) Inverted grayscale FISH images showing *slc6a3^+^* cells in maintained tail after knockdown of *soxB1-2* compared to control. D) Quantification of cell number in three 10000 µm^2^ boxes. The average of each box is plotted on the graph. Unpaired t-test. E) FISH images showing *laminin* expression in regenerated pharynges 9 dpa after knockdown of *foxA* compared to control. (**** P < 0.0001). Scale bars = 50 µm

**Extended Figure 10:**
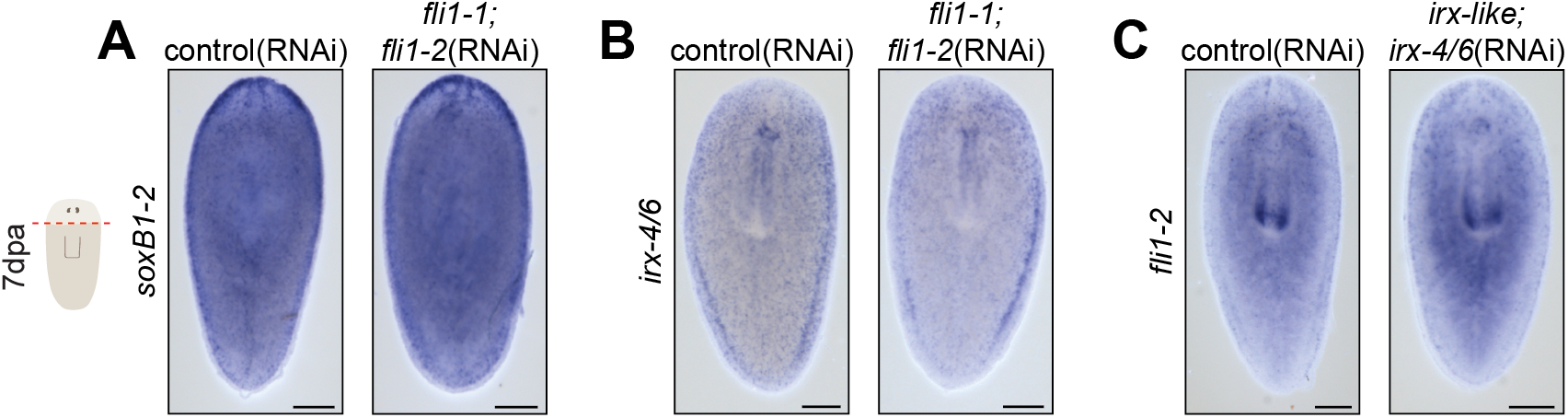
Transcription factor-encoding genes regulate each other. A) Whole-body colorimetric ISH showing *soxB1-2* expression 7 days post head amputation after double knockdown of *fli1-1;fli1-2.* B) Whole-body colorimetric ISH showing *irx-4/6* expression 7 days post head amputation after double knockdown of *fli1-1;fli1-2.* C) Whole-body colorimetric ISH showing *fli1-2* expression 7 days post head amputation after double knockdown of *irx-like;irx- 4/6.* Scale bars = 200 µm

---

**Supplemental File 1:**
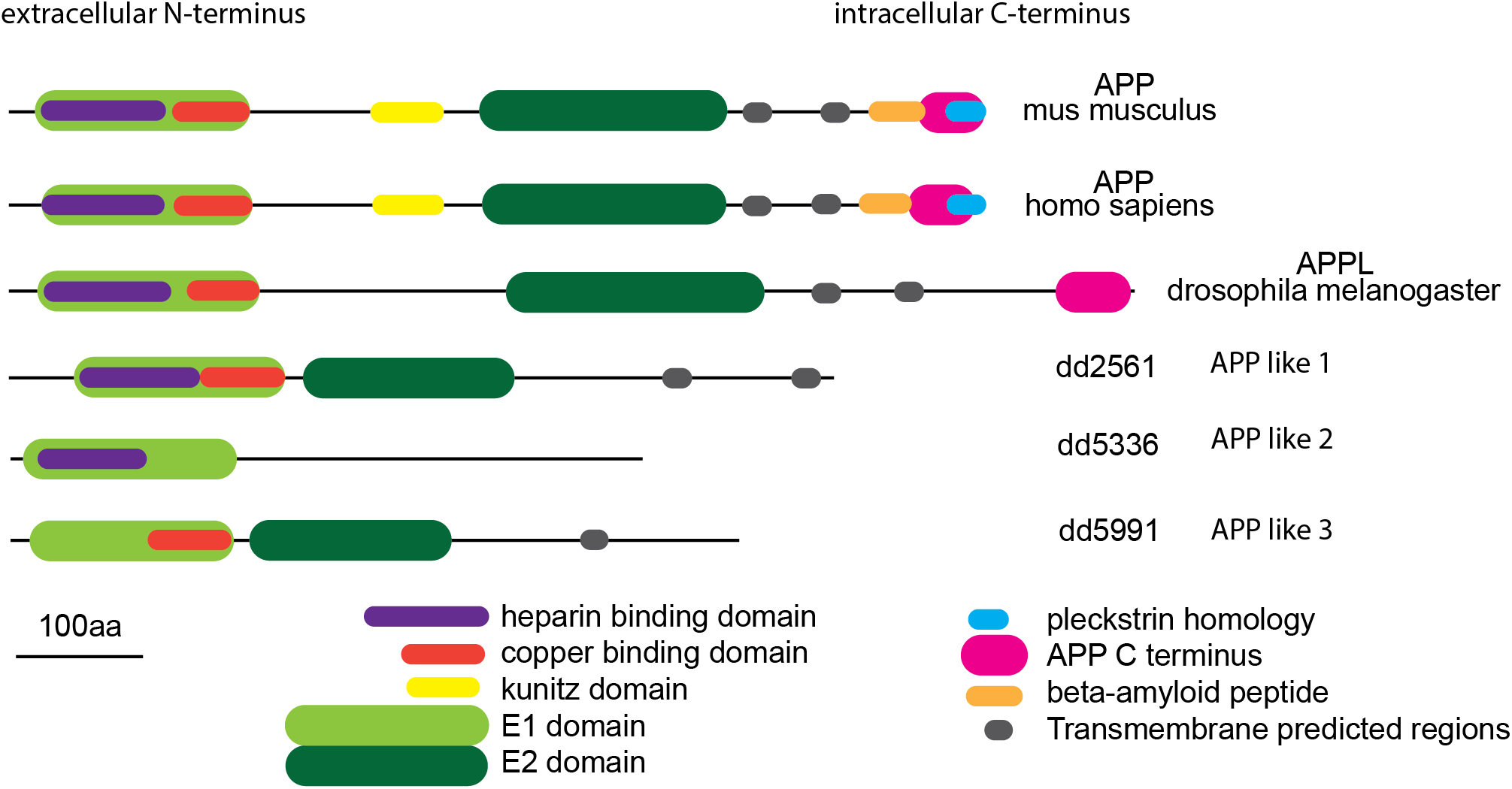
App phylogeny.

**Supplemental File 2.**
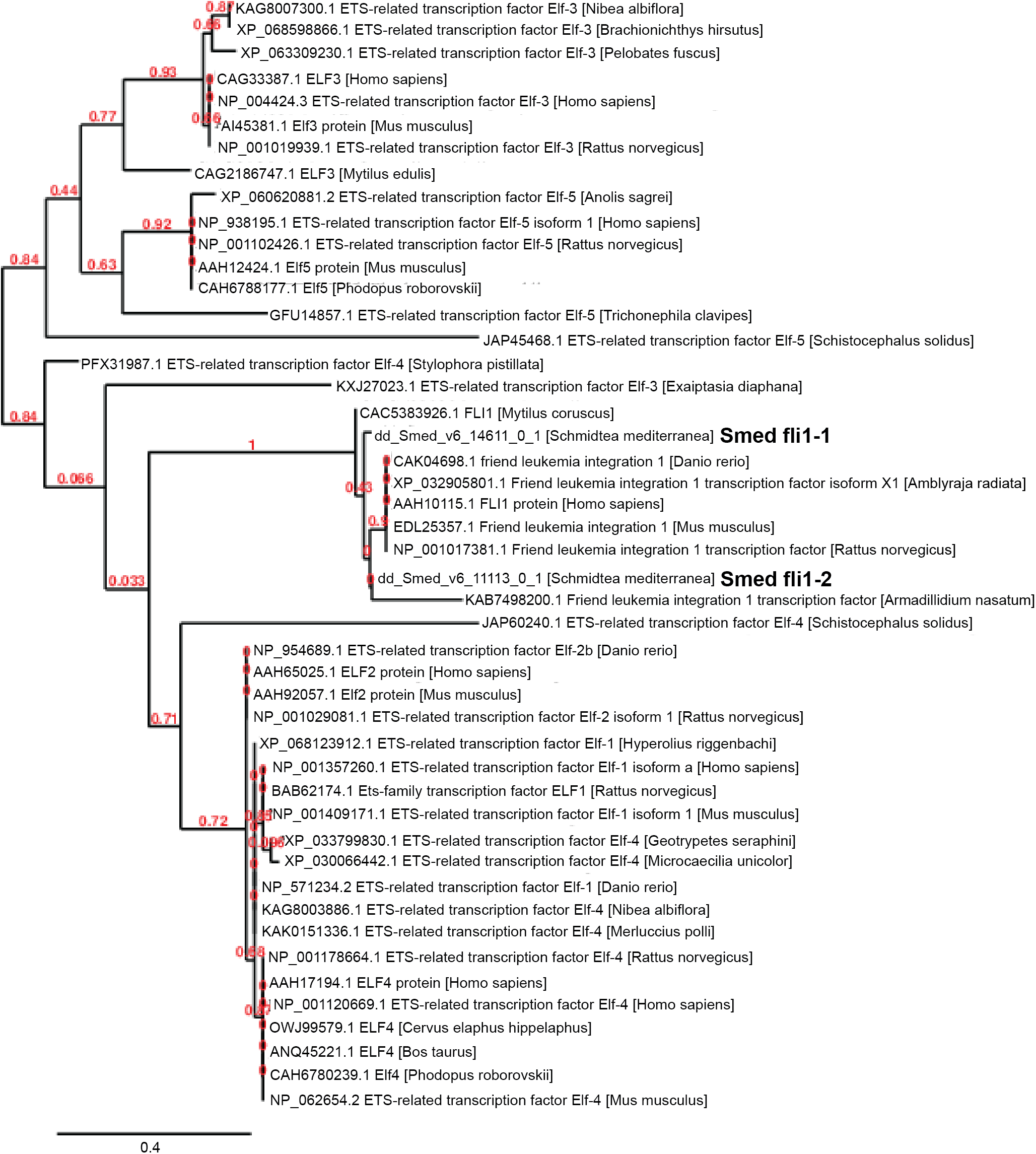

**Supplemental File 3.**
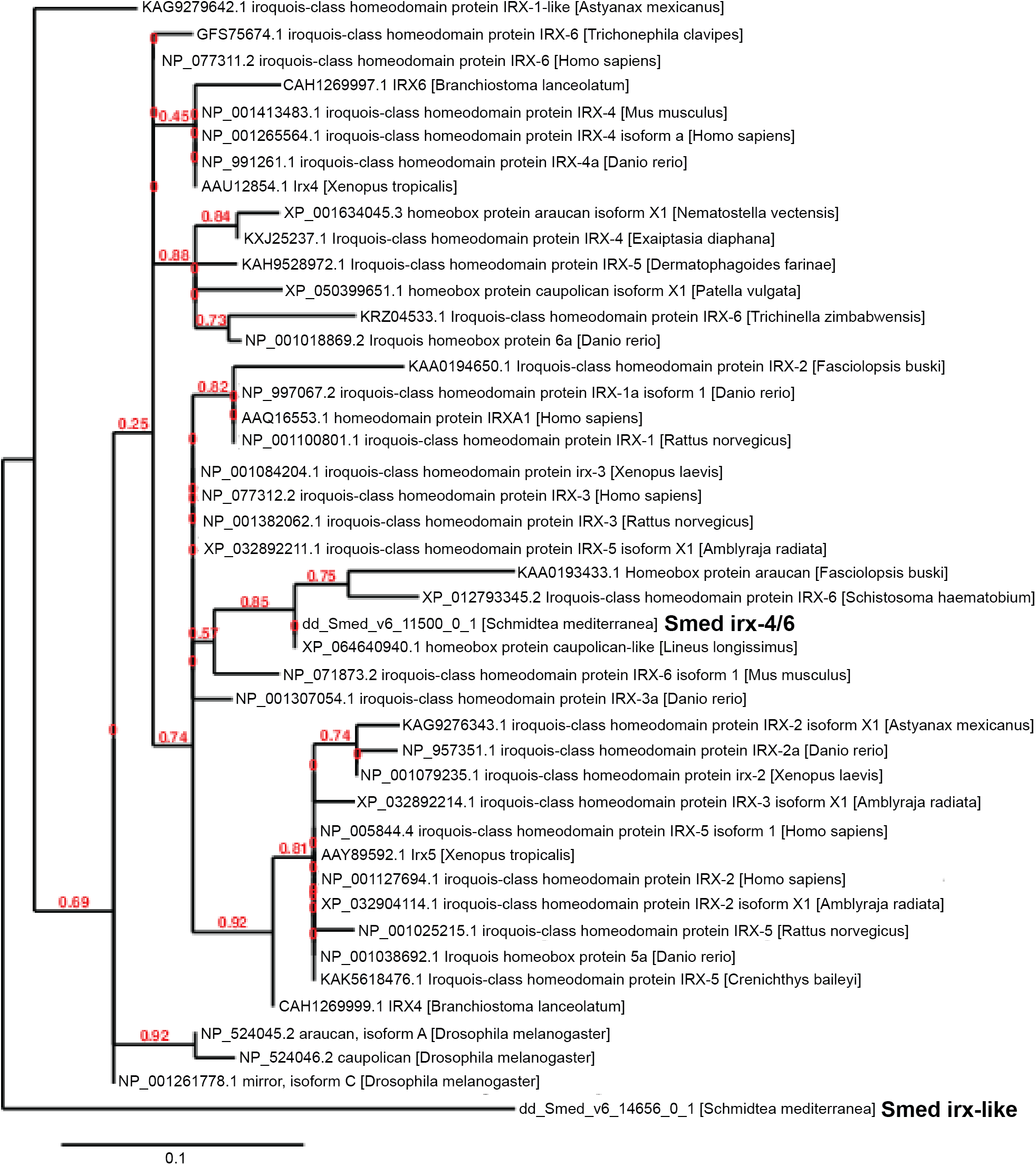

